# AdipoR2 is Essential for Membrane Lipid Homeostasis in Response to Dietary Saturated Fats

**DOI:** 10.1101/2020.06.11.144329

**Authors:** Ranjan Devkota, Mario Ruiz, Henrik Palmgren, Marcus Ståhlman, Himjyot Jaiswal, Marcello Maresca, Mohammad Bohlooly-Y, Xiao-Rong Peng, Jan Borén, Marc Pilon

**Affiliations:** Dept. Chemistry and Molecular Biology, Univ. Gothenburg, 405 30 Gothenburg, Sweden; Metabolism Bioscience, Early CVRM, BioPharmaceuticals R&D, AstraZeneca, Gothenburg, 431 83 Mölndal, Sweden; Dept. Molecular and Clinical Medicine/Wallenberg Laboratory, Institute of Medicine, Univ. of Gothenburg, 405 30 Gothenburg, Sweden; Discovery Biology, Discovery Sciences, R&D, AstraZeneca, Gothenburg, 431 83 Mölndal, Sweden; CellinkAB, Arvid Wallgrens Backe 20, 413 46 Gothenburg, Sweden

## Abstract

Membrane lipid composition influences vital processes in all types of cells. The mechanisms by which cells maintain membrane lipid homeostasis while obtaining most of their constituent fatty acids from a varied diet remain largely unknown. In an attempt to discover such mechanisms, we performed an unbiased forward genetic screen in *Caenorhabditis elegans* and conclude that the adiponectin receptor 2 (AdipoR2) pathway is essential to prevent saturated fat-mediated cellular toxicity. Transcriptomics, lipidomics and membrane property analyses in human HEK293 cells and primary human endothelial cells further support our conclusion that the essential function of AdipoR2 is to respond to membrane rigidification by promoting fatty acid desaturation. Our results demonstrate that AdipoR2-dependent regulation of membrane homeostasis is a fundamental mechanism conserved from nematodes to mammals that prevents saturated fat-mediated lipotoxicity.

**ONE SENTENCE SUMMARY:** The AdipoR2 protein insures membrane homeostasis in response to dietary saturated fatty acids that promote membrane rigidification.

## INTRODUCTION

Cellular membrane lipid composition is constantly adapting to meet the demands of essential functions such as cell division, growth, secretion or signaling. On the other hand, dietary manipulations readily influence membrane composition (*1, 2*), and therefore variations in diets pose different challenges for membrane homeostasis. In particular, an excess of dietary SFAs, such as palmitic acid (PA, C16:0) (Fig. S1A), is detrimental to cell physiology and induces lipotoxicity (*3*). Conversely, diets rich in monounsaturated fatty acids (MUFAs), such as oleic acid (OA, C18:1) and polyunsaturated fatty acids (PUFAs) such as eicosapentaenoic acid (EPA, C20:5) (Fig. S1A), have protective effects (*4*). The mechanisms governing membrane homeostasis in response to dietary variations are still poorly understood compared to other processes, such as glucose or cholesterol homeostasis; here, we seek to identify and elucidate such mechanisms.

## RESULTS

### Forward genetics identifies PAQR-2 and IGLR-2 as essential for SFA tolerance

Using *C. elegans*, we performed the first whole-animal forward genetic screen to identify genes that are required for the ability to tolerate dietary SFAs (Fig. 1A). This screen led to the isolation of five novel mutants that were completely intolerant of dietary SFAs (Fig. 1B, C). When these mutants were given dietary PA, their membrane phospholipids had abnormally high levels of SFAs (Fig. 1D) and reduced levels of MUFAs and PUFAs (Fig. S1B, C). These changes were accompanied by pronounced membrane rigidification measured *in vivo* using a fluorescence recovery after photobleaching (FRAP) assay (Fig. 1E).

**Fig. 1.**
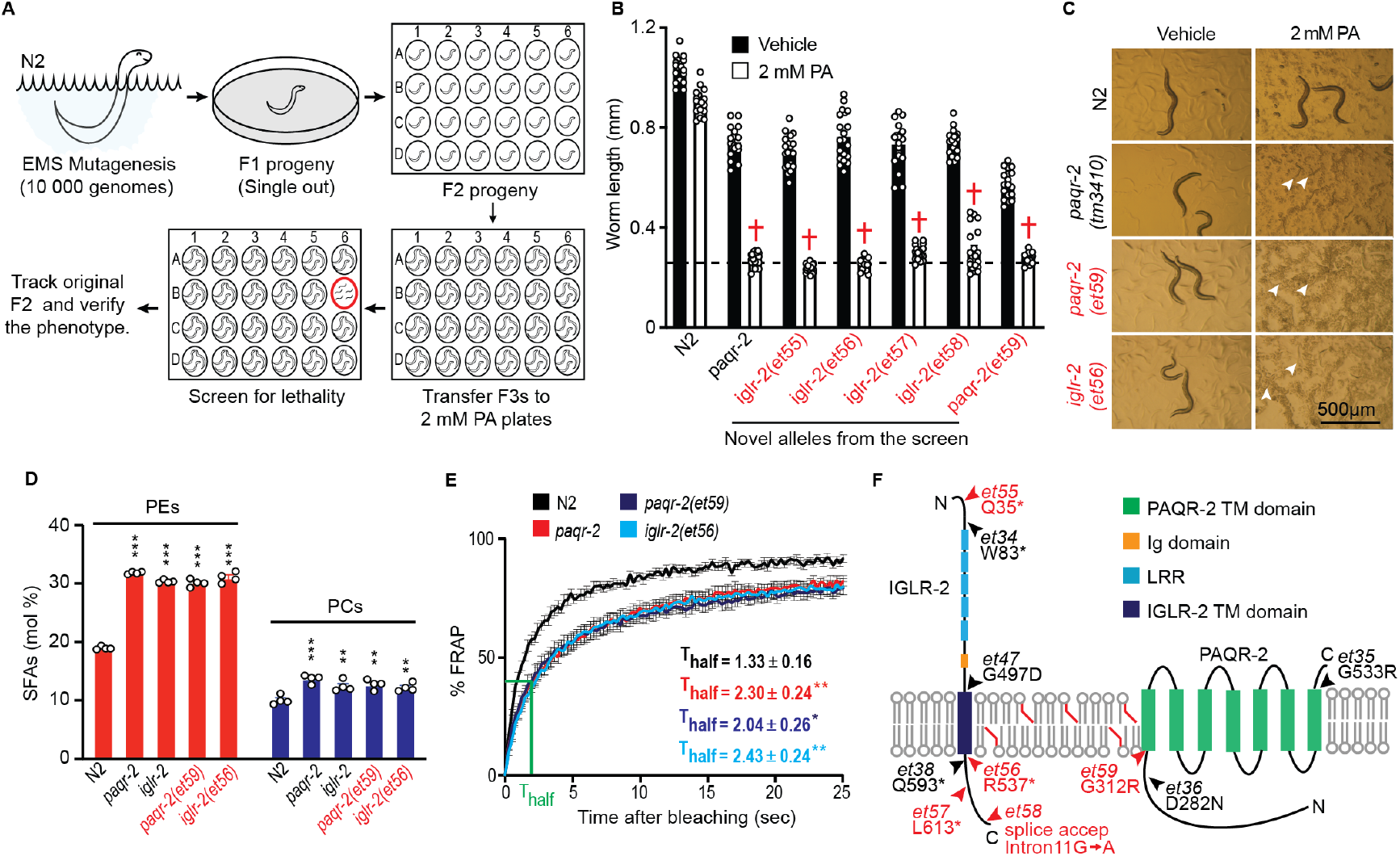
Screen for PA-sensitive mutants identifies only novel alleles of *paqr-2* and *iglr-2* in *C. elegans*. **A**, Screening strategy to isolate mutants sensitive to PA. **B**, Growth assay of five novel mutants (red text) in vehicle vs. 2 mM PA treatment (n=20 for all genotypes and treatments). The dashed line represents the average length of L1 worms at the start of the experiment. † symbolises lethality of the mutant worms treated with PA. **C**, Representative pictures of PA-sensitive mutants. Arrowheads point towards the dead worms. **D**, SFAs abundance (mol%) in phosphatidylethanolamines (PEs) and phosphatidylcholines (PCs) in worms treated with 2 mM PA (n=4 for all genotypes). **E**, Membrane rigidity in worms treated with 2 mM PA quantified by half-time of recovery in FRAP experiments (T_half_ value) (N2, n=8; *paqr-2*, n=6; *paqr-2(et59)*, n=7; *iglr-2(et56)*, n=7). **F**, The isolated mutants are novel alleles (in red) of *paqr-2* and *iglr-2*; previously available alleles (in black) are also indicated. *paqr-2* or *iglr-2* refer to the reference null alleles *tm3410* and *et34*, respectively.

On the basis of genetic complementation and genomic DNA sequencing, we determined that four out of the five mutants were loss-of-function (*lof)* alleles of IGLR-2, a homolog of mammalian LRIG-type proteins, and that the fifth mutant was a *lof* allele of PAQR-2, a homolog of the mammalian adiponectin receptors (AdipoRs) (Fig. 1F). The identity of the mutations was further confirmed by genetic rescue with wild-type transgenes (Fig. S1D, E). Published structural studies of the human AdipoRs suggest that they act as ceramidases (*5, 6*). Remarkably, previous studies have shown that the primary cellular function of PAQR-2 and AdipoR2 is to promote fatty acid desaturation upon membrane rigidification and hence regulate membrane homeostasis (*7, 8*), that PAQR-2 forms a complex with IGLR-2, and that these two proteins are mutually dependent for their function in membrane homeostasis (*9*). As expected, the identified mutant alleles of *paqr-2* and *iglr-2* were able to maintain membrane fluidity in normal conditions (Fig. S1F) but were sensitive to membrane rigidifying conditions such as cultivation in the presence of glucose, which is readily converted to SFAs by the *E. coli* that constitutes the diet of the worms *(7)*, and cold, which rigidifies membranes thermodynamically (Fig. S1G, H). Additionally, the novel mutants displayed the abnormal tail tip phenotype characteristic of *paqr-2* and *iglr-2* mutants (Fig. S1I). Based on these results, we conclude that *paqr-2* and *iglr-2* are essential for the ability of *C. elegans* to tolerate dietary SFAs. No other mutations were isolated in our PA-sensitivity screen, though our approach excludes essential genes which could not be isolated since they would cause lethality even in the absence of PA.

### Membrane rigidification promotes the PAQR-2/IGLR-2 interaction

The essential roles of PAQR-2 and IGLR-2 in preventing membrane rigidification suggest that they may be regulated by membrane properties. To test this possibility, we used the fluorescence resonance energy transfer (FRET) method (Fig. 2A). We found that the interaction between PAQR-2 and IGLR-2 in the gonad sheath cell membrane, where they are most abundantly expressed (*9*), was highly dynamic and that the strength of their interaction correlated with membrane rigidity across a range of conditions (Fig. 2B-E). In a negative control experiment, FRET did not detect any interaction between IGLR-2 and PAQR-1, a homolog of the mammalian adiponectin receptor 1 that does not interact with IGLR-2 (*9*) (Fig. S2A-C). We conclude that the formation of the PAQR-2/IGLR-2 complex is influenced by membrane properties and could therefore act as a plasma membrane fluidity sensor in eukaryotes analogous to membrane homeostasis regulators such as IRE1 in the endoplasmic reticulum (ER)(*10*), PCYT1 in the inner nuclear membrane (*11*) and DesK in bacterial membranes (*12*).

**Fig. 2.**
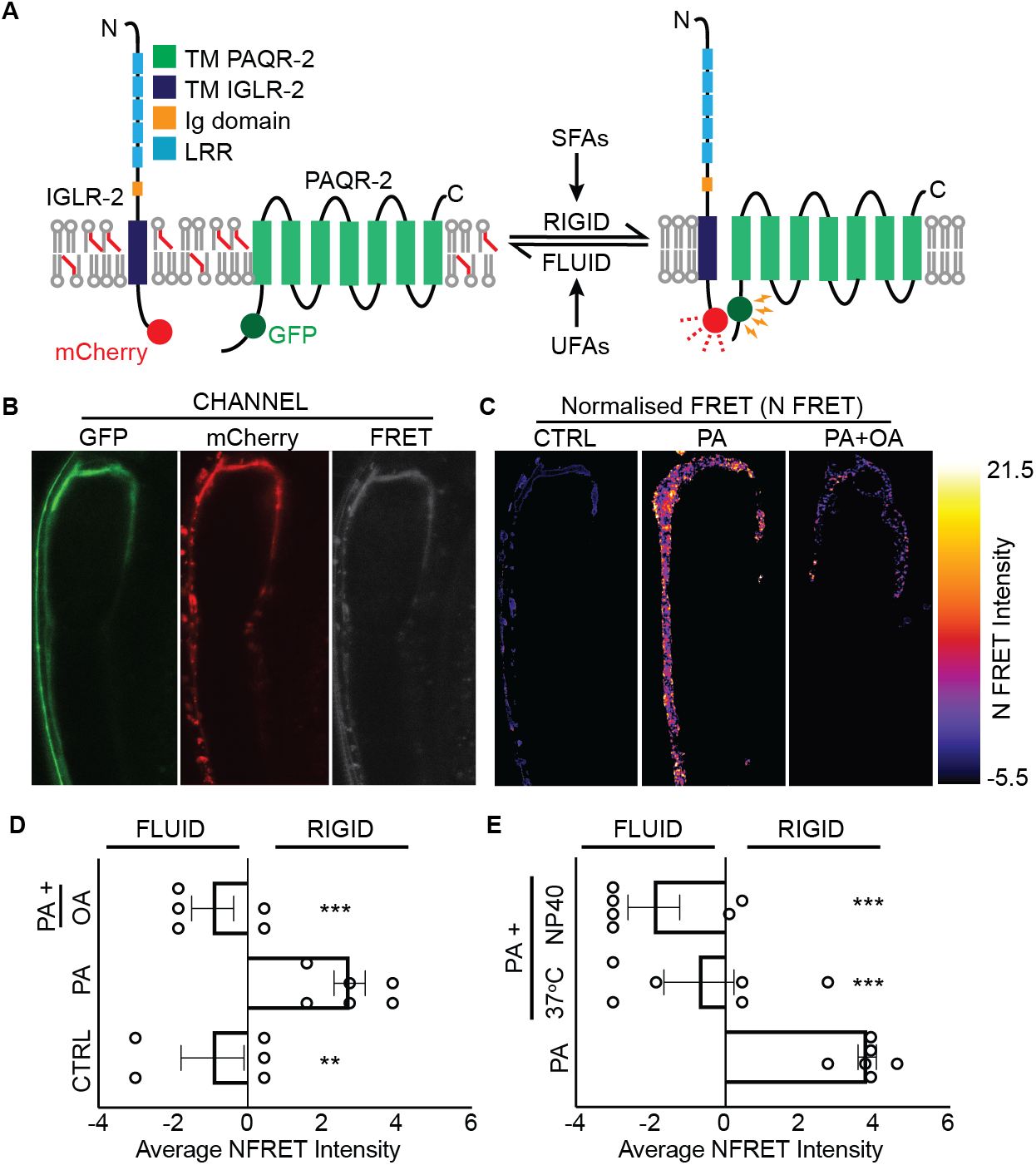
PAQR-2/IGLR-2 act as fluidity sensors. **A**, Model of PAQR-2/IGLR-2 interaction influenced by membrane fluidity and monitored by FRET. **B**, PAQR-2∷GFP and IGLR-2∷mCherry expression in gonad sheath cells as indicated in the GFP and mCherry detection channels. The distinct FRET signal between PAQR-2∷GFP and IGLR-2∷mCherry is shown in the FRET channel. **C**, PixFRET-based normalised FRET signals under different treatments. **D, E**, Average normalized FRET intensity in membrane rigidifying or fluidizing conditions; NP40 is Nonidet P-40, a mild detergent (CTRL, n=5; 2 mM PA, n=6; 2 mM PA+ 2 mM OA, n=5; 2 mM PA+37°C, n=6; 2 mM PA+ 0.1% NP40, n=6).

### SFAs cause membrane rigidification in AdipoR2-deficient cells

To investigate conservation with mammalian physiology, we generated AdipoR2 knockout (KO) HEK293T cells using CRISPR/Cas9 (Fig. S3A) and performed a detailed phenotypic characterization. AdipoR2-KO cells showed delayed growth in basal condition and, importantly, were extremely sensitive to PA supplementation (Fig. S3B, C). This sensitivity could be reversed by the addition of low concentrations of the membrane-fluidizing EPA (Fig. S3D). Subsequently, we measured membrane order using the laurdan dye generalized polarization (GP) index and FRAP, and found that the cellular membranes of PA-challenged AdipoR2-KO cells were markedly packed, indicating increased rigidity (Fig. 3A, B, Fig. S3E-H). This response to PA was again reversed by addition of EPA (Fig. S3I, J). Consistently, PA-treated AdipoR2-KO cells had excessive levels of SFAs and depletion of MUFAs and PUFAs in phosphatidylcholines (PCs) and phosphatidylethanolamines (PEs), and these defects increased with time of PA treatment (Fig. 3C, D) (Fig. S4A-D). Separately, we used atomic force microscopy (AFM), which can measure mechanical stiffness at atomic scales, to verify membrane rigidification in AdipoR2-knockdown HEK293 cells (Fig. S5A-C), and also found that the membranes of AdipoR2-knockdown cells became excessively rigid when challenged with other SFAs, i.e. myristic acid (C14:0) or stearic acid (C18:0), and thus is not a PA-specific phenomenon (Fig. S5D). Taken together, these findings indicate that AdipoR2-dependent regulation of membrane homeostasis is essential for the ability of human cells to tolerate saturated fats. It should be noted that AdipoR2 was initially discovered as a putative adiponectin receptor (*13*). However, none of our experiments included adiponectin supplements and the membrane homeostasis functions of AdipoR2 therefore appears to be adiponectin independent, which is consistent with a previous study (*8*).

**Fig. 3.**
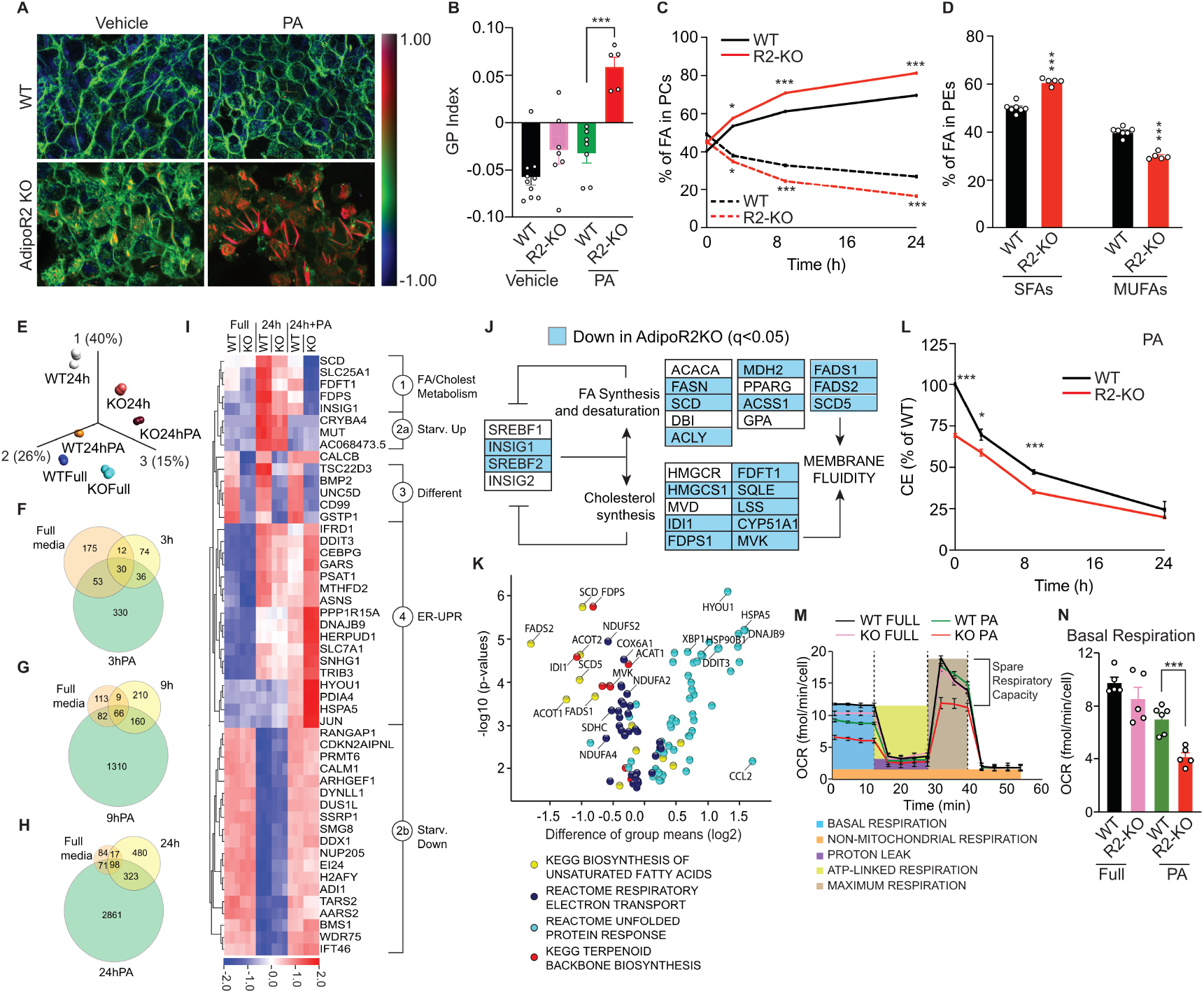
AdipoR2 is important for normal cellular response to PA. **A**, Representative pseudocolor images of laurdan dye GP index for WT and AdipoR2-KO HEK293 cells challenged with 200 μM PA. **B**, Average GP index for WT and AdipoR2-KO cells for different treatments (WT Vehicle, n=10; AdipoR2-KO Vehicle, n=7; WT PA, n=7; AdipoR2-KO PA, n=5;). **C**, SFA (solid lines) and MUFA (dashed lines) abundance (mol%) in PCs in WT and AdipoR2-KO cells challenged with 200 μM PA for 0, 3, 9 and 24 h (WT 0 h, n=3; WT 3 h, n=3; WT 9 h, n=3; WT 24 h, n=7; R2-KO n=5 for all timepoints). **D**, SFA and MUFA abundance (mol%) in PEs in WT and AdipoR2-KO cells challenged with 200 μM PA for 24 h (WT 24 h, n=7; R2-KO n=5). **E**, Principal component analysis for three condition (full medium, 24 h starvation in serum-free medium and 24 h in serum-free medium containing 200 μM PA) for WT and AdipoR2-KO cells. **F-H**, Venn diagrams showing the number of genes for which the expression was significantly different (q<0.05) between WT and AdipoR2-KO cells under each condition and time point. **I**, Hierarchically ordered heat map of the 50 genes showing the most significant variation (p≤4.5e-26) in a multigroup comparison among full medium, 24 h serum-free medium and 24 h serum-free medium containing 200 μM PA. **J**, Simplified view of the SREBP pathway that influences membrane composition, with genes downregulated in the AdipoR2-KO cells after 24 h on PA highlighted in blue. **K**, Volcano plot of genes from four selected pathways and with q<0.05 when comparing WT and AdipoR2 KO cells after 24 h in the presence of PA. **L**, Relative levels of cholesteryl esters (CEs) in WT and AdipoR2-KO cells when challenged with 200 μM PA for 0, 3, 9 and 24 h (WT 0 h, n=3; WT 3 h, n=3; WT 9 h, n=3; WT 24 h, n=7; R2-KO n=5 for all timepoints). **M**, Seahorse based oxygen consumption rates (OCR) in WT and AdipoR2-KO cells in full medium or in serum-free medium containing 200 μM PA. **N**, Basal respiration in WT and AdipoR2-KO cells under the indicated treatments (WT Full medium, n=5; WT serum-free medium + 200 μM PA, n=6; AdipoR2-KO Full medium, n=6; AdipoR2-KO serum-free medium + 200 μM PA, n=5).

### AdipoR2 is required for a normal cellular response to PA

To better understand the overall consequences of AdipoR2 deficiency, we performed a transcriptomic analysis of control and AdipoR2-KO HEK293T cells grown in full medium, serum-free media or serum-free media supplemented with 200 μM PA for 0, 3, 9 and 24 h. Principle component analysis of all monitored transcripts (over 50 000) showed clear transcriptome differences among the genotypes and treatments (Fig. 3E). Of note, the SCD gene, which encodes the human Δ9 desaturase, was the most significantly downregulated gene in the AdipoR2-KO cells after 24 h of PA treatment. Globally, the transcriptional response to PA in AdipoR2-KO was increasingly abnormal with time: 330 genes were mis-regulated after 3 h, 1 310 after 9 h and 2 861 after 24 h treatment with PA (Fig. 3F-H). We also identified 53 genes that were differently expressed (either up or down) in AdipoR2-KO cells in the presence of PA at all three time points tested (Fig. S6A), many of which are important for cholesterol synthesis (e.g. CYP51A1), fatty acid desaturation (FADS2), TCA cycle/respiration (e.g. IDH2, NDUFA6, NDUFB1) or are components of ribosomes (e.g. RPL36AL, RPL5, RPL6). Further, 17 genes were mis-regulated (either up or down) in the AdipoR2-KO cells under all conditions (Fig. S6B), which suggests that the loss of AdipoR2 led to a change in the differentiation state of these cells.

A volcano plot of all 3 050 genes with q < 0.05 and at least a 1.2-fold difference between control and AdipoR2-KO cells after 24 h with PA showed that 1 556 were upregulated and 1 494 were downregulated in the AdipoR2-KO cells (Fig. S6C). The top 50 genes showing the most significant variation among all treatments clustered into four different categories: 1) genes involved in fatty acid and cholesterol metabolism, 2) genes that responded to starvation, 3) genes that always differed between WT and AdipoR2-KO cells and 4) genes involved in the ER-unfolded protein response (UPR) (Fig. 3I, Fig. S6D). Importantly, we observed that the genes in the first category were strongly downregulated whereas genes in the fourth category were strongly upregulated in AdipoR2-KO cells upon PA treatment. This suggests that membrane homeostasis failure is the fundamental defect in AdipoR2-KO cells and is associated with an inability to regulate unsaturated fatty acid (UFA) and cholesterol synthesis in response to membrane rigidification; the observed ER stress response is most likely a consequence of the defective membrane homeostasis.

Sterol response element binding proteins (SREBPs) are known regulators of fatty acid synthesis/desaturation and cholesterol synthesis (*14*). We found that upon PA treatment, AdipoR2-KO cells transcriptionally phenocopied the loss of SREBPs (*15*), with most of the pathway being downregulated by 24 h (Fig. 3J). Using siRNA in human cells, we found that inhibition of the SREBFs (Fig. S7A) led to reduced SCD expression (Fig. S7B), lower membrane fluidity (Fig. S7C-G) and altered membrane lipid composition (Fig. S7H-N) in a manner analogous to AdipoR2-KO cells when challenged with PA. The observed downregulation of the SREBP pathway therefore likely contributes to the membrane phenotype of the AdipoR2-KO cells.

Gene set enrichment analysis (GSEA) (*16*) using the KEGG (*17*) and REACTOME (*18*) collections of gene sets showed that: (1) loss of AdipoR2 downregulated pathways involved in ribosomes, glycolysis/gluconeogenesis and metabolism of xenobiotics by cytochrome P450 under all conditions; and (2) PA treatment of AdipoR2-KO cells led to the downregulation of pathways involved in propanoate metabolism, oxidative phosphorylation and steroid biosynthesis (Fig. S8A-C). Further, an analysis of all genes in several selected pathways supported the interpretation that a primary membrane homeostasis defects in AdipoR2-KO cells led to an increased ER stress response: pathways involved in membrane homeostasis (biosynthesis of unsaturated fatty acids, terpenoid backbone biosynthesis) and respiratory electron transport were downregulated while genes of the ER-UPR pathway were upregulated (Fig. 3K, Fig. S8D-J). Detection of SCD and FADS2 using western blots showed that their protein levels concur with the results of the transcriptomic analysis, i.e. were decreased in the AdipoR2-KO cells challenged with PA (Fig. S9A-C). Consistent with the downregulated transcripts of genes involved in cholesterol synthesis and electron transport chain, the relative levels of cholesteryl esters and mitochondrial respiration were both reduced in AdipoR2-KO cells challenged with PA (Fig. 3L-N, Fig. S9D-H). Several of our key results were independently verified using an independently derived AdipoR2-KO clone (Fig. S10). Altogether, our findings show that AdipoR2 is essential for the transcriptional response required for membrane homeostasis when cells are challenged with SFAs.

### AdipoR2 prevents SFA toxicity

Others have previously shown that genes such as SCD, FADS2, ACSL4 and PEMT, which are important for fatty acid desaturation and incorporation of unsaturated lipids into cellular membranes, are crucial for protecting cells against PA toxicity (*19, 20*). To investigate how AdipoR2 compares with these other genes, we efficiently silenced AdipoR2, SCD, FADS2, ACSL4 and PEMT in HEK293 cells (Fig. S11A). A time-course FRAP experiment showed that as early as 6 h after PA treatment, and persisting over time, inhibition of AdipoR2, SCD or FADS2 caused the strongest rigidification of cellular membranes (Fig. 4A, Fig. S11B-I). This result was independently confirmed using the laurdan dye method (Fig. 4B-C, Fig. S11J). Importantly, results from a comparative lipid analysis [that measured SFAs, MUFAs and PUFAs in PCs and PEs, PC/PE ratio, relative levels of lysophosphatidylcholine (LPC), free cholesterol (FC), total triacylglycerides (TAGs) and TAG 48:0] showed that silencing AdipoR2 or SCD has the most profound effect on lipid composition, followed by silencing of FADS2, ACSL4 or PEMT, which had only a weak effect (Fig. 4D-F, Fig. S11K-R). Further, and as expected, we observed that membrane rigidification was always accompanied by increased ER-UPR levels (Fig. 4G, Fig. S11S-U). Of note, AdipoR2 ranked 4^th^ and 25^th^ in two recently published genome-wide CRISPR/Cas9 screens to identify genes required to prevent PA toxicity or desaturase inhibition (by low oxygen) toxicity (*20, 21*). We therefore conclude that AdipoR2 is one of the most important genes to not only sense toxic membrane rigidification caused by elevated extracellular SFAs, but also to trigger an adaptive response that normalizes membrane lipid composition.

**Fig. 4.**
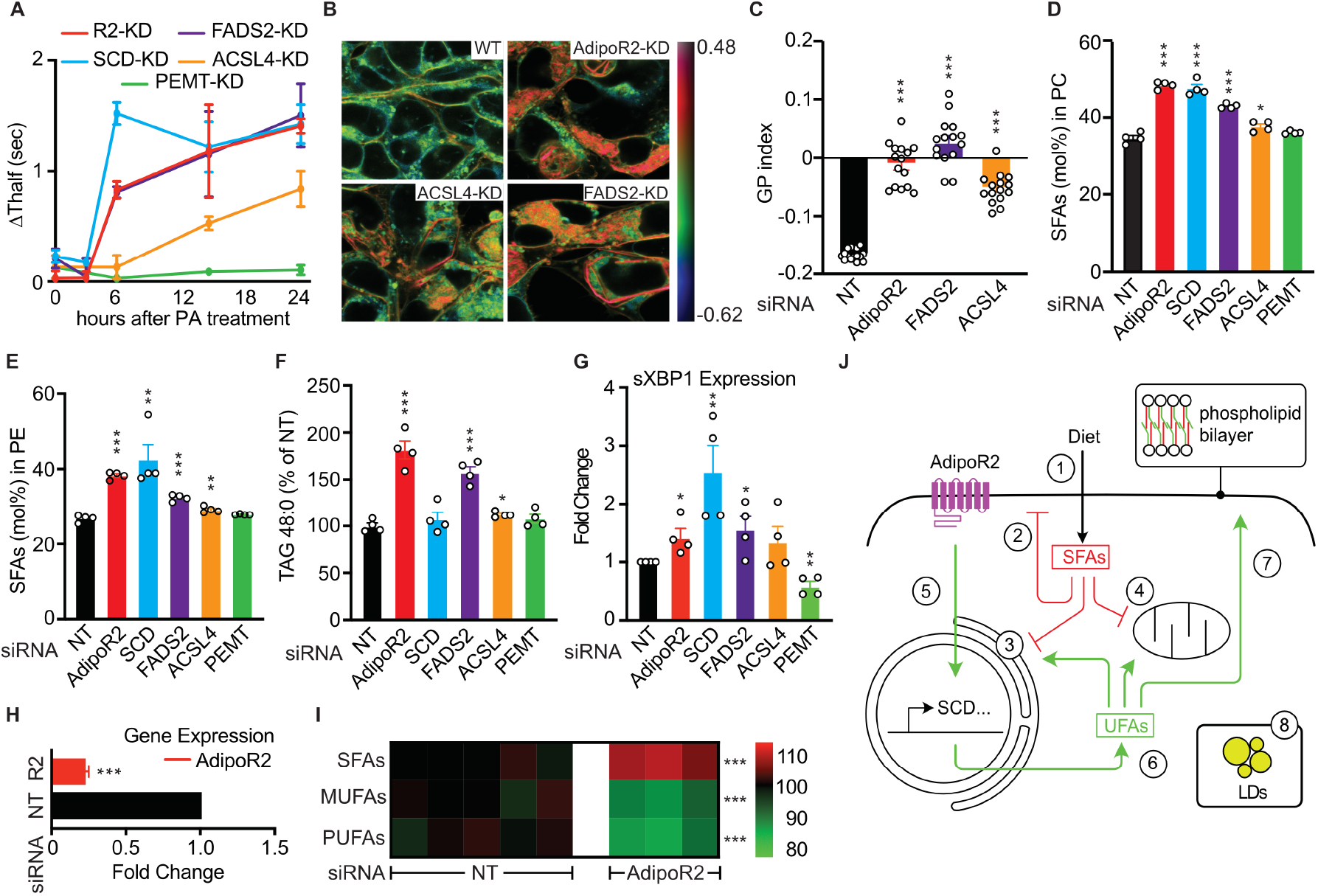
AdipoR2 regulates membrane homeostasis to counter PA-induced lipotoxicity. **A**, Differential T_half_ values comparing AdipoR2, SCD, FADS2, ACSL4 and PEMT knockdown (KD) in HEK293 cells challenged with 200 μM PA in a time-course FRAP experiment. **B**, Representative pseudocolor images and **C**, average laurdan dye GP index in HEK293 cells treated with AdipoR2, ACSL4 or FADS2 siRNA and challenged with 200 μM PA (n=15 for all treatments). **D**, **E** and **F,** SFAs abundance (mol%) in PCs and PEs and relative levels of TAG 48:0 in HEK293 cells with the indicated genes knocked down and challenged with 200 μM PA (n=4 for all treatments). **G**, Spliced XBP1 mRNA expression in HEK293 cells with the indicated genes knocked down and challenged with 200 μM PA (n=4 for all treatments). **H**, Efficiency of AdipoR2 silencing in HUVECs measured using qPCR. **I**, Relative levels of SFAs, MUFAs and PUFAs in PCs in HUVECs treated with NT (nontarget) and AdipoR2 siRNA and challenged with 200 μM PA for 6 h (NT siRNA, n=5; AdipoR2 siRNA, n=3). **J**, AdipoR2-dependent membrane regulation to counter SFA-induced lipotoxicity: Dietary SFAs are taken up by the cells and become incorporated into membrane phospholipids (1), leading to plasma membrane rigidification (2), ER stress (3), and impairment of mitochondria function (4). In healthy cells, AdipoR2 responds to increased membrane rigidification by signalling to promote the expression of desaturases and other lipid metabolism genes (5), leading to increased levels of UFAs available for incorporation into phospholipids (6) and normalization of membrane fluidity (7). Some cells can also sequester SFAs into lipid droplets in the form of TAGs (8). A similar process occurs in *C. elegans*, where rigidification activates the AdipoR homolog PAQR-2 by promoting its interaction with IGLR-2.

Finally, we found that silencing AdipoR2 in primary human umbilical vein endothelial cells (HUVECs) (Fig. 4H) led to the accumulation of membrane-rigidifying SFAs and depletion of membrane-fluidizing MUFAs and PUFAs when the cells were treated with PA (Fig. 4I). AdipoR2 is therefore required to maintain membrane homeostasis in human primary cells.

### Conclusions

Excessive levels of SFAs lead to cellular dysfunction and death, a phenomenon that has been established in a variety of cells including endothelial cells, fibroblasts, pancreatic β cells, hepatocytes and cardiomyocytes (*22*). Here, we performed a genetic screen to understand the molecular basis of SFA-mediated lipotoxicity and found that AdipoR2-dependent regulation of membrane homeostasis is crucial for cells to tolerate SFAs. Our further analyses indicate that excess SFAs in the cells results in SFA-rich membrane phospholipids leading to membrane rigidification, ER stress, and mitochondrial dysfunction. We propose that a healthy cell responds to membrane rigidification via AdipoR2-dependent induction of fatty acid desaturases and other lipid metabolism genes; this response leads to increased levels of MUFAs and PUFAs available for incorporation into phospholipids and thereby normalizes membrane fluidity and cellular processes (Fig. 4L). In conclusion, we identified an evolutionarily conserved membrane homeostatic pathway essential for cells to sense toxic membrane rigidification caused by elevated extracellular SFAs and mount an adaptive response; these findings should prove valuable in many disease contexts where lipotoxicity or membrane composition defects have been implicated.

## MATERIALS AND METHODS

### *C. elegans* Strains, Transgenes and Cultivation

The wild-type *C. elegans* reference strain N2 and the mutant alleles studied (except for the novel alleles from this study) are available from the *C. elegans* Genetics Center (CGC; USA). The *pPAQR-2∷GFP* construct used for transgenic rescue experiments and as a template for construction of other plasmids has been described elsewhere (*23*). Unless otherwise stated, experiments using the *C. elegans* strains maintenance and experiments were performed at 20°C using the *E. coli* strain OP50 as food source. *E. coli* was maintained on LB plates kept at 4°C (re-streaked every 6-8 weeks); single colonies from these plates were cultivated overnight at 37°C in LB medium, then used for seeding NGM plates (*24*). OP50 stocks were kept frozen at −80°C and new LB plates were streaked every 3-4 months. NGM plates containing 20 mM glucose were prepared using a 1 M stock solution that was filter sterilized then added to the molten NGM medium.

### Pre-loading of *E. coli* with PA

A stock of 0.1 M PA (Sigma) was dissolved in ethanol and diluted in LB medium to a final concentration of 2 mM, inoculated with OP50 bacteria, then shaken overnight at 37°C. The bacteria were then washed twice with M9 to remove traces of PA and growth medium, diluted to equal concentrations based on optical density (OD_600_), concentrated 10X by centrifugation, dissolved in M9 and seeded onto NGM plates lacking peptone (200 μl/plate). Worms were added the following day.

### Screen for PA intolerance

N2 worms were mutagenized for 4 h by incubation in the presence of 0.05 M ethyl methane sulfonate (EMS) according to the standard protocol (*24*). The worms were then washed and placed on a culture dish. After 2 h, vigorous L4 hermaphrodite animals were transferred to new culture plates. Four days later, individual worms from F1 progenies were transferred to separate culture plates and, after a further four days, 4 F2s from each F1 plate were transferred onto 24-well plates. After three days, F3s were transferred to new 24-well plates containing OP50 pre-loaded with 2 mM PA. Four days later, the plates were screened for lethality and phenotype was further verified with original F2 plates.

### Genetic complementation and sequencing

Genetic complementation was performed as described elsewhere(*25*). The coding sequences of *paqr-2* and *iglr-2* novel alleles were amplified using different sets of primers. The following primers were used to cover the entire coding sequence of *paqr-2*: 5’-tccgctttattctctcacag-3’ and 5’-gaattagcggtaactgaccact-3’; 5’-atcactgcttgccctcatat-3’ and 5’-gagatcggtgtaaatgatgcga-3’; 5’-tcgcatcatttacaccgatctc-3’ and 5’-caatagctcacatgtgtgca-3’; 5’-tgcacacatgtgagctattg-3’ and 5’-ccaaatatcgcactttccag-3’; 5’-gataaattcgcggagcccaaa-3’ and 5’-agcagcttcaagaagagcat-3’. The following primers were used to cover the coding sequence of *iglr-2*: 5’-atcctcgaactttccatcgt-3’ and 5’-ccattcagagattccaaatcacc-3’; 5’-gcaccatttccgcaactagaa-3’ and 5’-cgttagatatcgagcagagaac-3’; 5’-atccttacgccgaaactcaa-3’ and 5’-atgctaaaatgtgtgcgcct-3’; 5’-aggcgcacacattttagcat-3’ and 5’-ttagaccgccgtaatcat-3’; 5’-atgattacggcggtctaa-3’ and 5’-cacataggacattttcga-3’; 5’-tcgaaaatgtcctatgtg-3’ and 5’-atgatgatggcaccgatt-3’.

### Construction of pIGLR-2∷mCherry

The pIGLR-2∷mCherry construct was generated with a Gibson assembly cloning kit (NEB) with the following four fragments: (1) *iglr-2* 3’ UTR and vector backbone was amplified from the *pIGLR-2* rescue construct(*9*) as a template using the primers 5’-aattctttaaatcattttgtttgtttgtctataaccattccaaaatcggtgcca-3’ and 5’-acgtcagggcgaattccagcacactggcggccgttactagtggat-3’, (2) *iglr-2* promoter was amplified from *pIGLR-2* rescue construct as a template using the primers 5’-ccagtgtgctggaattcgccctgacgtgacctacgtccctattttgca-3’ and 5’-aaaatacaaattttcgcatttcttcttttctttgtatcaagacatctgcgatgt-3’, (3) *iglr-2* gene was amplified from *pIGLR-2* rescue construct as a template using the primers 5’-aaagaaaagaagaaatgcgaaaatttgtatttttcgtcgtagctattcttattca-3’ and 5’-taataattgccatgttatcttcttcaccctttgagaccattctcttttctggtggagaatct-3’, (4) mCherry was amplified using the primers 5’-atggtctcaaagggtgaagaagataacatggcaattattaaagagtttatgcgtttcaaggt-3’ and 5’-ggttatagacaaacaaacaaaatgatttaaagaattacttatacaattcatccatgcca-3’.

### Construction of PAQR-2p∷PAQR-1∷GFP

The PAQR-2p∷PAQR-1∷GFP construct was generated with a Gibson assembly cloning kit (NEB) with the following three fragments: (1) *paqr-2* promoter with the vector backbone was amplified from *pPAQR-2∷GFP* construct using the primers 5’-aagggcgaattctgcagatatcca-3’ and 5’-ctcctttactcattgatgccattttgttaaagctgaattt-3’, (2) *paqr-1* gene with 3’UTR was amplified using N2 worm lysate as a template using the primers 5’-tggcggttctggtggcggatctaatccagatgaggtcaatcg-3’ and 5’-tctgcagaattcgcccttttttactcttaatatccgag-3’, (3) GFP was amplified from *pPAQR-2∷GFP* construct using the primers 5’-ggcatcaatgagtaaaggagaagaacttttcactggagtt-3’ and 5’-ccgccaccagaaccgccacctttgtatagttcatccatgc-3’.

### Generation of transgenic *C. elegans*

Plasmids were prepared with ThermoFisher GeneJet plasmid miniprep kit and used with the following concentrations: test plasmids at 25 ng/μl, *pPD118.33* (*Pmyo-2∷GFP*) (Addgene plasmid #1596) at 5 ng/μl and *pBSKS* (Stratagene) at 70 ng/μl for a total of 100 ng/μl. *pPAQR-2∷GFP and pIGLR-2∷mCherry* co-injection was done with concentration of 25 ng/μl for each of the plasmid.

### Growth and tail tip scoring assays

For length measurement studies, synchronized L1s were plated onto test plates seeded with *E. coli*, and worms were mounted and photographed 144 h (15°C experiments) or 72 h (all other experiments) later. The length of 20 worms were measured using ImageJ (*26*). Quantification of the withered tail tip phenotype was done on synchronous 1-day old adult populations cultivated at 20°C, that is 72 h post L1 (n ≥100).

### FRET

FRET experiments were carried out on strains *pPAQR-2∷GFP* (donor alone), *pIGLR-2∷mCherry* (acceptor alone) and *pPAQR-2∷GFP; pIGLR-2∷mCherry* (donor and acceptor). The worms were bleach synchronized and after 48 h, they were washed and transferred onto: (1) non-peptone control plates, (2) non-peptone plates containing OP50, pre-loaded with 2 mM PA, or (3) non-peptone plates containing OP50 pre-loaded with 2 mM PA and 2 mM OA. For detergent rescue experiments, worms were transferred to 0.1% NP40 non-peptone plates containing OP50 pre-loaded with 2 mM PA. After 24 h, individual day 1 adult worms were placed on microscopic glass slide and immobilized using 10 mM levamisole. For 37°C experiments, worms treated with OP50 pre-loaded with PA were incubated in the live cell chamber of the microscope for 30-45 mins. FRET images were acquired with an LSM880 confocal microscope equipped with a live cell chamber and ZEN software (Zeiss) with a 40X water immersion objective. For GFP channel, 488 nm laser with 2% laser intensity was used for excitation and the emission was recorded between 494 and 565 nm. For the mCherry channel, 561 nm laser with 0.2% laser intensity was used for excitation and the emission was recorded between 588 and 696 nm. For the FRET channel, 488 nm laser with 2% laser intensity was used for excitation and the emission was recorded between 588 and 696 nm. All the images were acquired with 16-bits image depth and 1068 × 1068 resolution using a pixel dwell of 1.97 μs. Images were analyzed using ImageJ with PixFRET plugin (*27*).

### Cell cultivation

HEK293 and HEK293T cells were grown in DMEM containing glucose 1 g/l, pyruvate and GlutaMAX and supplemented with 10% fetal bovine serum, 1% non-essential amino acids, HEPES 10 mM and 1% penicillin and streptomycin (all from Life Technologies) at 37°C in a water-humidified 5% CO_2_ incubator. Cells were sub-cultured twice a week at 90% confluence. HUVECs were obtained from Gibco and cultivated as described(*8*). Briefly, cells were grown in M200 medium (Gibco) containing Low Serum Growth Supplement (Gibco) and 1% penicillin and streptomycin. Cells were sub-cultured twice a week at 90% confluence, and cultivated on treated plastic flasks and multi-dish plates (Nunc). For FRAP experiments, cells were seeded in glass-bottom dishes (Ibidi) pre-coated with 0.1% porcine gelatin (Sigma). For laurdan dye experiments, HEK293 cells and HUVECs were seeded in gelatin coated glass-bottom dishes (Ibidi) and HEK293T cells in CellCarrier 384-well plates (PerkinElmer). For atomic force microscopy experiments, cells were seeded in glass bottom dishes (WillCo Wells) pre-coated with 0.1% porcine gelatin. The HEK293 cell line was authenticated by genotyping and confirmed to be free of mycoplasma.

### Shotgun lipidomics in *C. elegans* and human cells

For worm lipidomics, samples comprised synchronized L4 larvae (one 9 cm diameter plate/ sample; each treatment/genotype was prepared in five independently grown replicates) grown overnight on OP50-seeded NGM or NGM containing 20 mM glucose. Worms were washed three times with M9, pelleted and stored at −80°C until analysis. For human cell lipidomics, cells (prepared in at least three independent replicates) were cultivated in the presence of 400 mM PA for 24 h prior to harvesting using TrypLE Express (Gibco). For lipid extraction, the pellet was sonicated for 10 min in methanol and then extracted according to published methods (*28*). Internal standards were added during the extraction. Lipid extracts were evaporated and reconstituted in chloroform:methanol [1:2] with 5 mM ammonium acetate. This solution was infused directly (shotgun approach) into a QTRAP 5500 mass spectrometer (Sciex) equipped with a with a TriVersa NanoMate (Advion Bioscience) as described previously (*29*). Phospholipids and cholesteryl esters were measured using precursor ion scanning (*30–32*) and TAGs were measured using neutral loss scanning (*33*). Free cholesterol from cells was quantified using straight phase HPLC coupled to ELS detection according to previous publication (*34*). The data were evaluated using the LipidView software (Sciex), and the complete lipid composition data are provided in Table S1.

### AdipoR2-KO Cell Line Generation and Sequencing

Genetical inactivation of the AdipoR2 gene was done in HEK293T cells using dual sgRNA transfection with Cas9 as described elsewhere (*35*). sgRNA sequences were: 5’-cgagccaacagaaaaccgat-3′ and 5′-caactggatggtacacgaag-3′. Individual cells were seeded and expanded. AdipoR2 genome-edition was characterized in two independent clones by Sanger sequencing.

### siRNA treatment

The following pre-designed siRNAs were purchased from Dharmacon: ACSL4 J-009364-05 AdipoR2 J-007801–10, FADS2 J-008211-09, Non-target (NT) D-001810–10, PEMT J-010392-05 SCD J-005061-07, SREBF1 J-006891-05 and SREBF2 J-009549-05. For HEK293 cells, transfection of 25 nM siRNA was performed in complete medium using Viromer Blue according to the manufacturer’s instructions 1X (Lipocalyx). HUVECs were transfected using 10 nM siRNA and Lipofectamine RNAiMAX Transfection Reagent following the HUVECs optimized protocol provided by the supplier (Invitrogen). Knockdown gene expression was verified 24 h (HUVECs) or 48 h (HEK293) after transfection.

### Quantitative PCR (qPCR)

Total cellular RNA was isolated using RNeasy Kit according to the manufacturer’s instructions (Qiagen) and quantified using a NanoDrop spectrophotometer (ND-1000; ThermoFisher). cDNA was obtained using a RevertAid H Minus First Strand cDNA Synthesis Kit with random hexamers (ThermoFisher). qPCR experiments were performed with a CFX Connect thermal cycler (Bio Rad) using Hot FIREpol EvaGreen qPCR SuperMix (Solis Biodyne) and standard primers. The relative expression of each gene was calculated according to the ΔΔCT method(*36*). Expression of the housekeeping gene PPIA (human samples) was used to normalize for variations in RNA input. Primers designed for this study were: PEMT-For (agcttctttgcactggggtt), PEMT-Rev (gggctggcgtgcatgat), SREBF1-For (gacctcgcagatccagcag), SREBF1-Rev (ataggcagcttctccgcatc), SREBF2-For (gtgctgttcctgactccctg) and SREBF2-Rev (cagccttcttcttggcctga). Primer sequences for ACSL4, AdipoR2, FADS2, PPIA, SCD were previously described(*34*), as were those for sXBP-1, ATF4, DDiT, HSPA5 (*37*).

### Cell culture fatty acid treatment

Myristic acid (14:0), PA (16:0) and stearic acid (18:0) were dissolved in sterile DMSO then mixed with fatty acid-free bovine serum albumin BSA (all from Sigma) in serum-free medium for 15 min at room temperature. The molecular ratio of BSA to fatty acid was 1 to 5.3 when using 400 μM of the SFA, and 1 to 2.65 when using 200 μM. Cells were then cultivated in this serum-free medium containing PA for 3, 6, 9 or 24 h prior to analysis. For some experiments EPA (20:5) (Sigma) was mixed with PA and together conjugated to BSA.

### FRAP in *C. elegans* and HEK293 cells

FRAP experiments in *C. elegans* were carried out using a membrane-associated prenylated GFP reported expressed on intestinal cells as described previously (*38*) and using a Zeiss LSM700inv laser scanning confocal microscope with a 40X water immersion objective. Briefly, the GFP positive membranes were photobleached over a circular area (seven-pixel radius) using 20 iterations of the 488 nm laser with 50% laser power transmission. Images were collected at a 12-bit intensity resolution over 256 × 256 pixels (digital zoom 4X) using a pixel dwell time of 1.58 μsec, and were all acquired under identical settings. For FRAP in mammalian cells, HEK293 and HEK293T cells were stained with BODIPY 500/510 C1, C12 (Invitrogen) at 2 μg/ml in PBS for 10 min at 37°C. FRAP images were acquired with an LSM880 confocal microscope equipped with a live cell chamber (set at 37°C and 5% CO_2_) with a 40X water-immersion objective as previously described (*8*). The recovery of fluorescence was traced for 25 s. Fluorescence recovery and T_half_ were calculated as described previously (*8*).

### Laurdan dye measurement of membrane order

Live HUVECs and HEK293 cells were stained with Laurdan dye (6-dodecanoyl-2-dimethylaminonaphthalene) (ThermoFisher) at 10 μM (HUVECs) or 15 μM (HEK293) for 45 min. Images were acquired with an LSM880 confocal microscope equipped with a live cell chamber (set at 37 °C and 5% CO_2_) and ZEN software (Zeiss) with a 40X water immersion objective as previously described (*8*). Cells were excited with a 405 nm laser and the emission recorded between 410 and 461 nm (ordered phase) and between 470 and 530 nm (disordered phase). For HEK293T, cells were stained with laurdan dye at 50 μM for 1 h. Images were captured using a robotic Yokogawa CV7000 spinning disc confocal microscope (Wako Automation). Temperature (37 °C) and carbon dioxide levels (5%) were controlled during live cell imaging. Images were acquired at 40X (NA.95, 2×2 binning) using a 405 nm laser excitation and 445/45 (ordered phase) plus 525/50 (disordered phase) bandpass filters using ZYLA 5.5 sCMOS cameras (Andor Technology). Pictures were acquired with 16 bits image depth and 1024 × 1024 resolution, using a pixel dwell of ~1.02 μs. Images were analyzed using ImageJ version 1.47 software, following published guidelines (*39*).

### Atomic force microscopy

Atomic force microscopy was carried out using a NanoScope V instrument (Bruker) and a Nikon Eclipse Ti-S inverted microscope. A pre-calibrated PeakForce QNM-Live Cell (PFQNM-LC) probe (Bruker AFM Probes) (tip length 17 μm, tip radius 65 nm) was used. The spring constant of the cantilever was 0.1010 N/m. Trigger threshold was set at 500 pm. Cells were in serum-free DMEM and at 37 °C during the experiment. NanoScope 8.15 software was used for data acquisition and NanoScope Analysis v1.5 for analysis (Bruker).

### Bioenergetics to assess mitochondrial function

Oxygen consumption rate (OCR) was determined by using a Seahorse XF 96 instrument (Agilent)(*40*). Briefly, cell medium composition during the assay was: Seahorse XF base medium (Agilent) supplemented with 5.5 mM glucose, 100 mM pyruvate and 2 mM L-glutamine. The Seahorse XF Cell Mito Stress Test was run using 2 μM oligomycin (Sigma-Aldrich), 200 μM DNP (as uncoupler; Sigma-Aldrich), 1 μM rotenone (Sigma-Aldrich) and 1 μM antimycin A (Sigma-Aldrich) as per the manufacturer’s instructions. Cells were stained with both Hoechst 33342 (ThermoFisher) and propidium iodide (ThermoFisher) to assess nuclei counts and viability as measured by fluorescence microscopy. OCR values were normalized against viable cells.

### IncuCyte growth assay

HEK293T cells were seeded at 5000 cells/well on clear 96 well culture plates (Corning). Cells were left to attach for 3 h. Four fields per well and at least five wells per treatment were imaged every 3 h using an IncuCyte™ S3 (Essen Bioscience) until cells reached confluency as determined by in-built confluency measurement algorithm. Once all conditions had reached confluency cells were washed twice with PBS before adding exogenous fatty acids in serum-free medium. Confluency was continuously assessed for 72 h post treatment.

### RNA sequencing and analysis

RNA was isolated using a RNeasy Mini Kit (Qiagen) with DNase treatment as per the manufacturer’s instructions. RNA concentration and quality was measured on a Fragment Analyser (Advanced Analytical Technologies). RNA integrity number greater than 9 was set as a quality control cut off for library preparation using 1 μg of total RNA. Poly(A) selected pair-end sequencing libraries were generated using the Illumina TrueSeq Stranded mRNA Sample Prep Kit (Illumina) according to the manufacturer’s instructions. Library quantification was assessed using the Standard Sensitivity NGS Kit (Advance Analytical Technologies) on a Fragment Analyser. Libraries were pooled and quantified on a Qubit Fluorometer using a DNA HS kit (ThermoFisher) followed by dilution to 1.8 pM before being sequenced using an Illumina NextSeq 500 Sequencer (Illumina). Sequencing reads were processed with the bcbio pipeline and mapped to the human genome (hg38). RNA-seq counts were generated using Salmon (*41*) which is an alignment free method that maps reads to human transcript sequences. Counts were imported into R using tximport (*42*) and collapsed to the gene level. The raw counts were adjusted using *DESeq2*, a method for differential analysis of count data, using shrinkage estimation for dispersions and fold changes to improve stability and interpretability of estimates (*43*). Normalized counts were imported into Qlucore Omics Explorer n.n (Qlucore AB) for analysis. Duplicates were “collapsed” to median values, resulting in a data set containing 54962 unique symbols (distinct transcripts) with read counts from each replicate. A threshold of 1 was applied to all entries and the counts were log2 transformed. The data were normalized for the purpose of heat map visualization (mean=0; variance=1). GSEA (*16*) for KEGG (*17*) and REACTOME (*18*) pathways v6.1 were done using two group comparisons where the KO24hPA treatment was compared to the other treatments using the following settings: min match set size=15, max match set size=500, permutations=100000, permutation method: variables and enrichment weight=1.

### Western blot for HEK293 cells

Protein lysates were prepared using RIPA buffer supplemented with 0.1% SDS, PhosSTOP and protease inhibitors (Sigma). Protein sample concentration was quantified using BCA protein assay kit (Pierce) according to manufacturer’s instructions. Equal amounts of protein were denatured in LDS sample buffer and sample reducing agent (Invitrogen) by heating to 70°C for 10 min. Samples were loaded on NuPAGE™ 4-12% Bis-Tris gels submerged in MOPS running buffer (Invitrogen) followed by transfer unto Invitrolon™ PVDF membranes (Invitrogen). All blots were run on the XCell SureLock mini-cell system (Invitrogen). Blots were blocked in 5% non-fat dry milk or BSA (Sigma) in PBS (Gibco) supplemented with 0.05% Tween-20 (Sigma) for 1 h at room temperature. Membranes were incubated overnight at 4°C with the primary antibody in manufacturer’s recommended blocking buffer. Blots were washed in PBS-tween and incubated with appropriate secondary antibody: goat anti-rabbit IgG/HRP (#7074, 1:3000, Cell Signaling) or goat anti-mouse IgG/HRP (1:5000, Dako) for 1 h at room temperature before being washed. Blots were developed using SuperSignal™ West Femto (ThermoFisher) and visualized using a ChemiDoc Touch MP imaging system (Bio-Rad). PageRule Plus (ThermoFisher) protein ladder was used to evaluate molecular weight. Blots were stripped for 15 min in Restore™ PLUS Western Blot Stripping Buffer (ThermoFisher) before blocking and proceeding to quantify GAPDH. Quantification was done using the Image Lab software normalizing the band intensity to that of GAPDH. Primary antibodies used were: mouse monoclonal anti-SCD1 (CD.E10, 1:1000, Abcam), rabbit polyclonal anti-FADS2 (#ab72189, 1:3000, Abcam), rabbit monoclonal anti-GAPDH (14C10, 1:1000, Cell Signaling).

### Statistics

Error bars show the standard error of the mean, and t-tests were used to identify significant differences between treatments. Asterisks are used in the figures to indicate various degrees of significance, where *: p<0.05; **: p<0.01; and ***: p<0.001.

## Supporting information

Table_S1_Lipidomics

## ACKNOWLEDGEMENTS

We acknowledge the Centre for Cellular Imaging at the University of Gothenburg and the National Microscopy Infrastructure for assistance with fluorescence microscopy, the Biochemical Imaging Centre Umeå for assistance with atomic force microscopy and the National Microscopy Infrastructure National Microscopy Infrastructure, NMI (VR-RFI 2016-00968). We also thank Fredrik Karlsson, Emma Svensk, Lisa Westlund, Matilda Colm and Rebecka Persson for their help with some of the experiments, and Rosie Perkins for comments on the manuscript. Funding: Cancerfonden, Swedish Research Council, Carl Tryggers Stiftelse, Diabetesfonden, Swedish Foundation for Strategic Research, Kungliga Vetenskaps-och Vitterhets-Samhället i Göteborg, Wilhelm och Martina Lundgrens Stiftelse, Längmanska Kulturfonden and Åke Wibergs Stiftelse.

## SUPPLEMENTARY MATERIALS

**Fig. S1.**
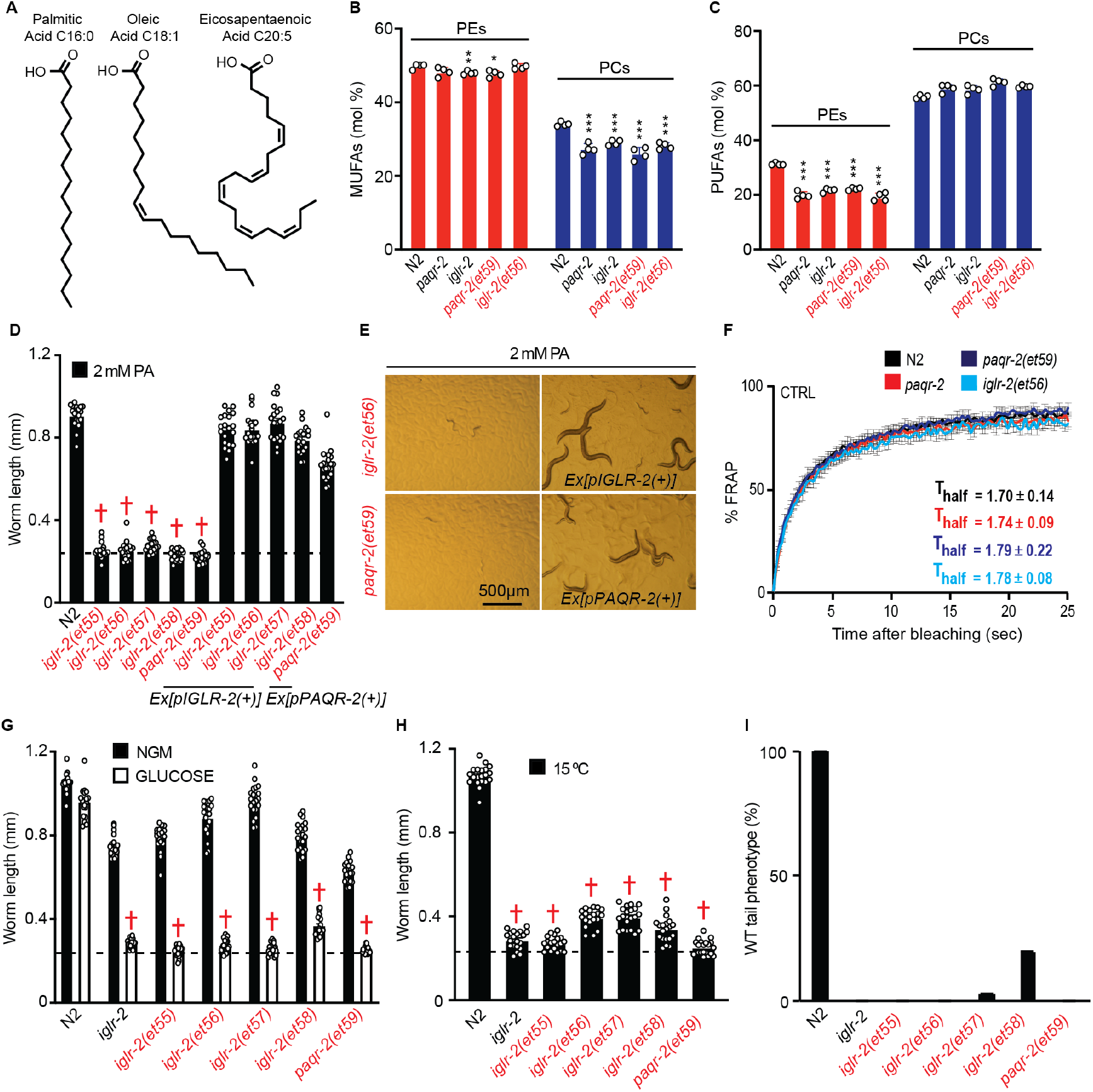
The novel *paqr-2* and *iglr-2* alleles are loss-of-function alleles. **A**, Illustration depicting different types of fatty acids: palmitic acid, oleic acid and eicosapentaenoic acid, examples of a SFA, MUFA and PUFA respectively. **B**, **C**, Abundance of MUFAs and PUFAs (mol%) in phosphatidylethanolamines (PEs) and phosphatidylcholines (PCs) in worms treated with 2 mM PA (n=4 for all genotypes; *paqr-2* and *iglr-2* refer to the reference null alleles *tm3410* and *et34*, respectively). **D**, Introduction of wild-type *paqr-2(+)* or *iglr-2(+)* transgene rescue the lethality of the novel mutants when challenged with 2 mM PA (n=20 for all genotypes). The dashed line represents the average length of L1 worms at the start of the experiment. † symbolizes of lethality. **E**, Representative pictures of the transgenic rescue of the mutants when treated with PA. **F**, Membrane fluidity measured using FRAP under control conditions (NGM plates) in WT and mutants (N2, n=3; *paqr-2*, n=5; *paqr-2(et59)*, n=5; *iglr-2(et56)*, n=3). **G**, **H**, Growth assay of the mutants in 20 mM glucose and 15°C (n=20 for all genotypes and treatments). **I**, Tail-tip phenotypes scored on 1-day-old adults (n=100 for all genotypes).

**Fig. S2.**
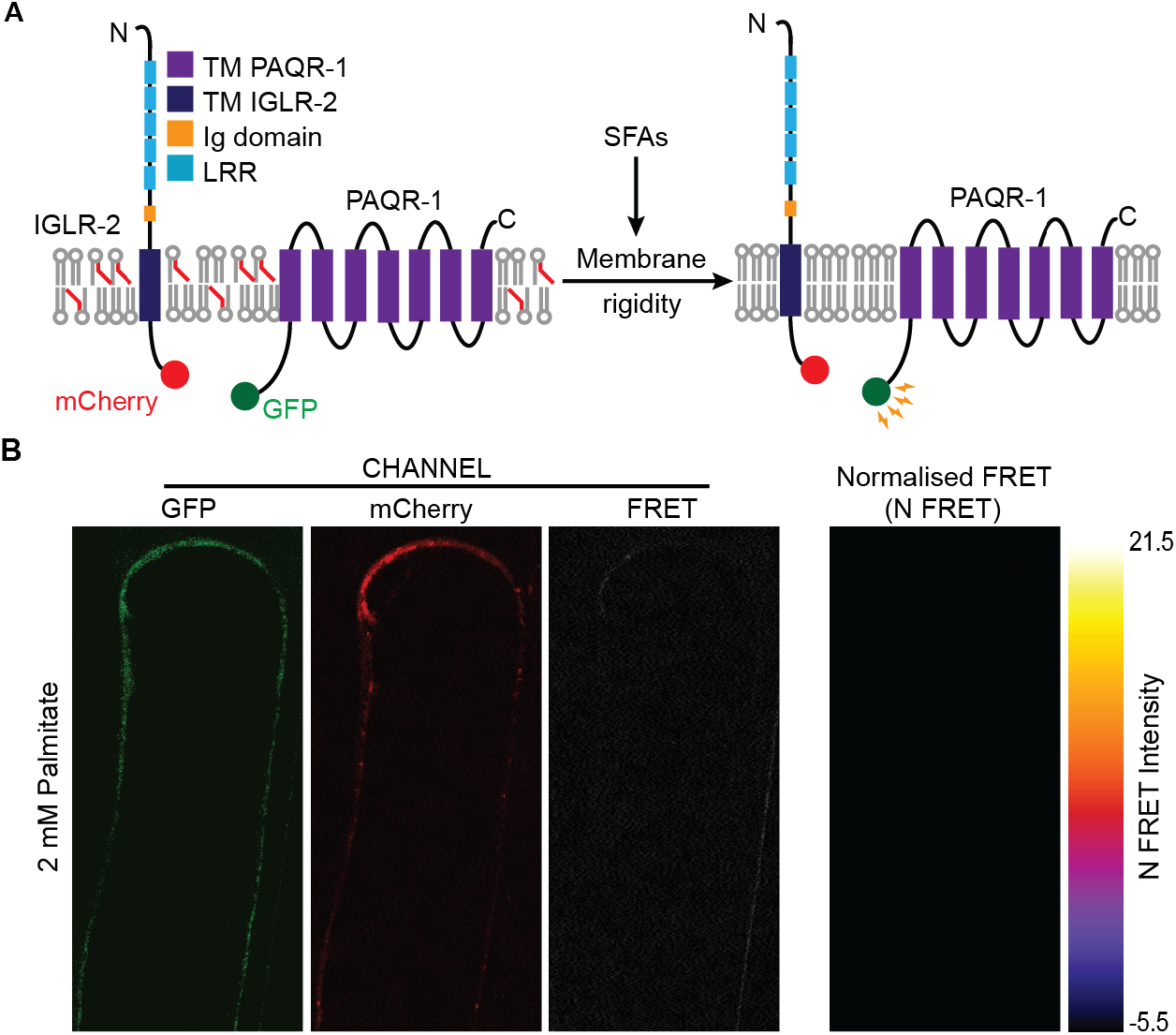
PAQR-1 does not interact with IGLR-2. **A**, Hypothetical model of interaction between PAQR-1 and IGLR-2. **B**, PAQR-1∷GFP and IGLR-2∷mCherry expression as indicated in the GFP and mCherry detection channels in gonad sheath cells from worms treated with 2 mM PA. Few background pixels in the FRET channel but absence of any signal upon PixFRET based normalization suggesting that PAQR-1 and IGLR-2 do not interact. This experiment is a negative control for Fig. 2.

**Fig. S3.**
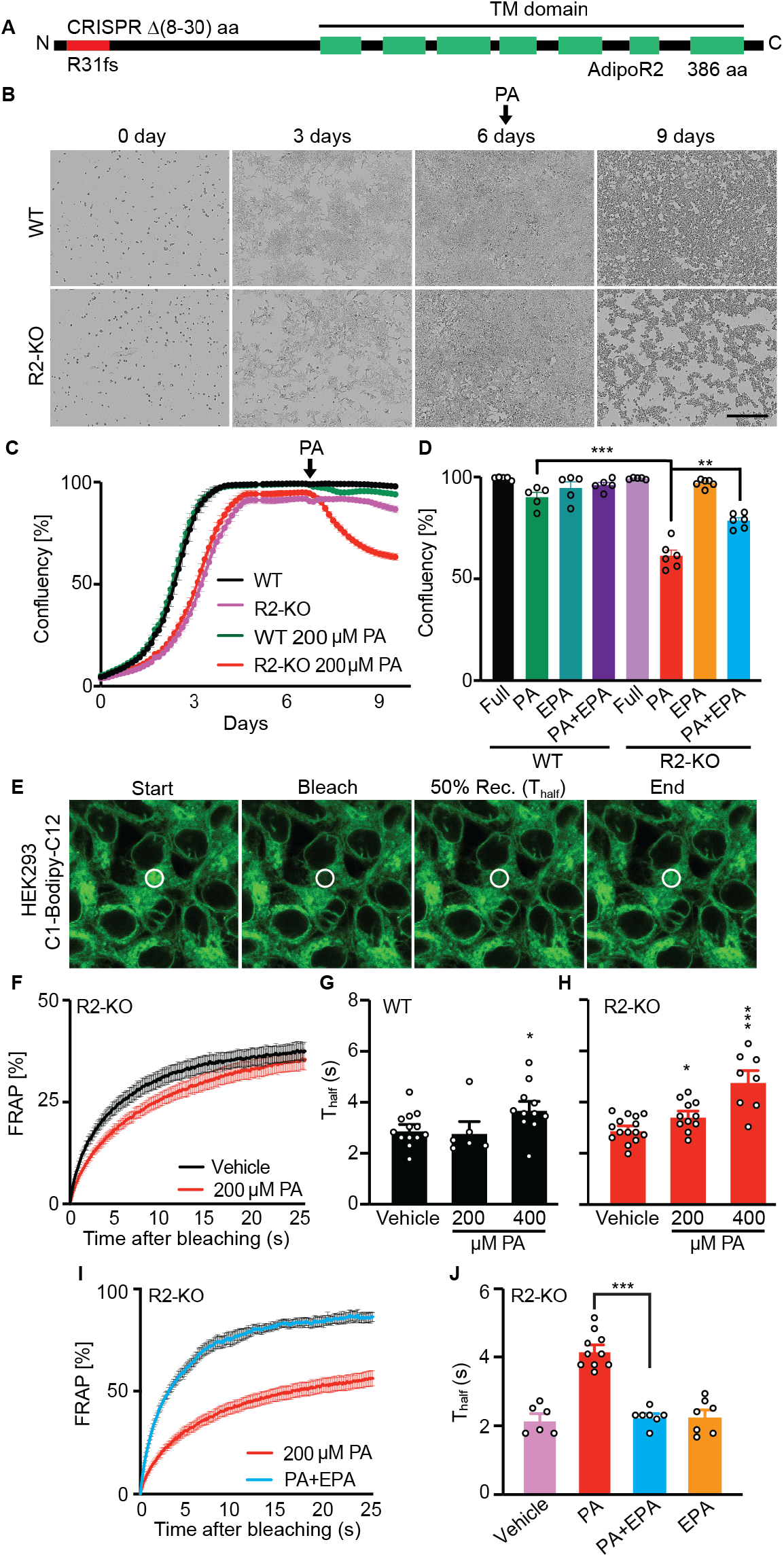
Characterization of HEK293 AdipoR2-KO cells. **A**, Representation of the AdipoR2 protein with the seven transmembrane domains indicated in green and the CRISPR-deleted amino acids (8-30) in red that result in a frameshift mutation at R31. **B**, Representative images indicating the morphology of WT and AdipoR2-KO cells pre- and post-PA treatment. **C**, Growth of WT and AdipoR2-KO cells pre- and post-PA treatment. (WT, n=6; AdipoR2-KO, n=4; WT PA, n=6; AdipoR2-KO PA, n=8). **D**, Growth of WT and AdipoR2-KO cells under different treatments (WT Full, n=5; WT PA, n=5; WT EPA, n=5; WT PA+EPA, n=5; AdipoR2-KO Full, n=5; AdipoR2-KO PA, n=6; AdipoR2-KO EPA, n=6; AdipoR2-KO PA+EPA, n=6). **E**, C1-Bodipy-C12 staining of HEK293T cells indicating different phases during a FRAP experiment (start, post-bleach, 50% recovery and end). The white circle represents the FRAP region. **F**, FRAP curves for AdipoR2-KO cells treated with vehicle or PA (AdipoR2-KO Vehicle, n=12; AdipoR2-KO PA, n=10). **G**, **H**, Average T_half_ values for WT and AdipoR2-KO cells treated with vehicle or PA (WT Vehicle, n=13; WT 200 μM PA, n=6; WT 400 μM PA, n=11; AdipoR2-KO Vehicle, n=15; AdipoR2-KO 200 μM PA, n=11; AdipoR2-KO 400 μM PA, n=8). **I**, FRAP curve for AdipoR2-KO cells treated with PA or PA and EPA (AdipoR2-KO PA, n=10; AdipoR2-KO PA+EPA, n=7). **J**, Average T_half_ values for AdipoR2-KO cells treated as indicated (AdipoR2-KO Vehicle, n=6; AdipoR2-KO PA, n=10; AdipoR2-KO PA+EPA, n=7; AdipoR2-KO EPA, n=7). PA and EPA were used at 200 μM and 20 μM, respectively.

**Fig. S4.**
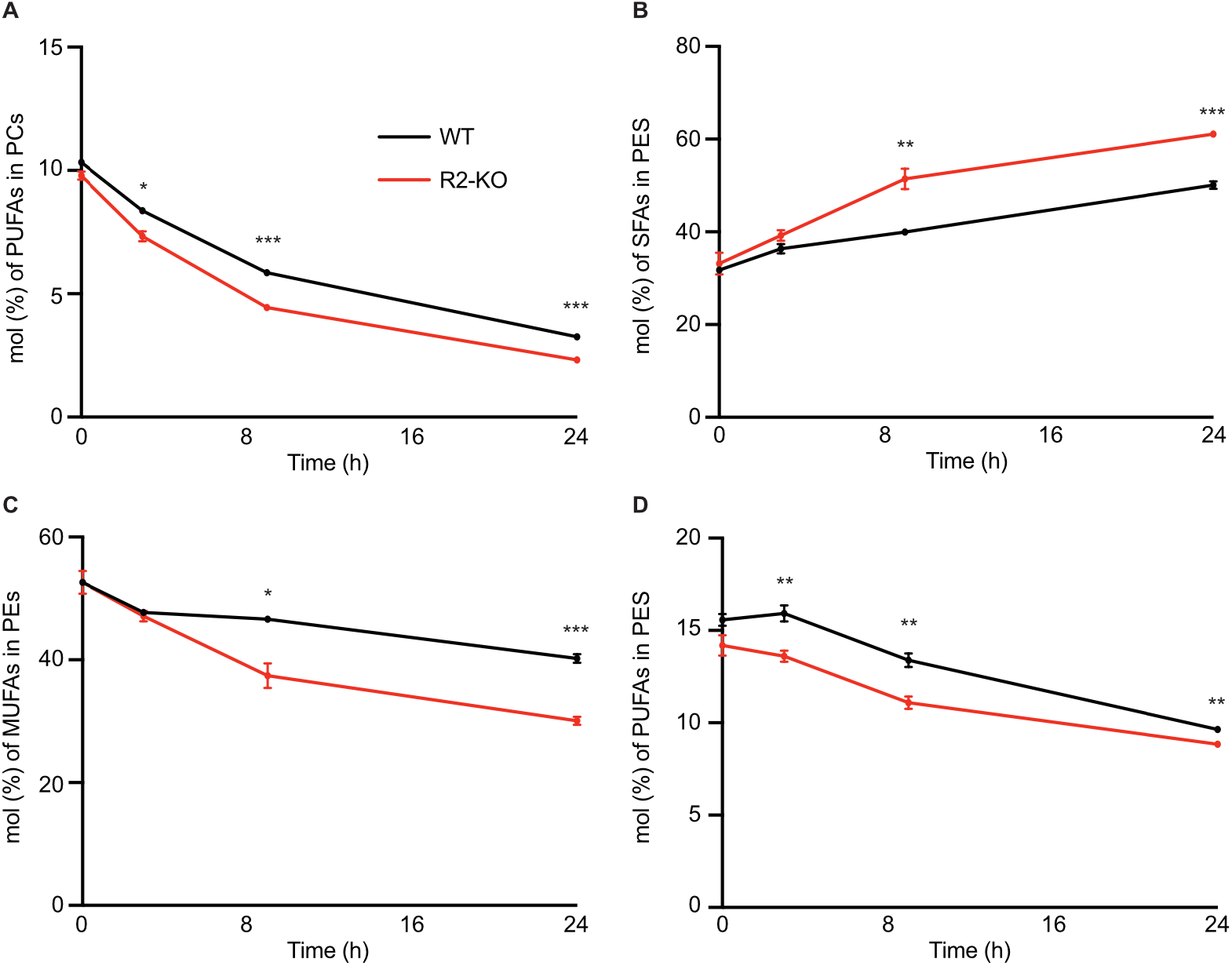
Time-course lipidomics in WT and AdipoR2-KO cells challenged with PA. **A**, PUFA abundance (mol%) in PCs in WT and AdipoR2-KO cells treated with 200 μM PA for 0, 3, 9 and 24 h (WT 0 h, n=3; WT 3 h, n=3; WT 9 h, n=3; WT 24 h, n=7; AdipoR2-KO n=5 for all timepoints). **B-D**, SFA, MUFA and PUFA abundance (mol%) in PEs in WT and AdipoR2-KO cells treated with 200 μM PA for 0, 3, 9 and 24 h (WT 0 h, n=3; WT 3 h, n=3; WT 9 h, n=3; WT 24 h, n=7; AdipoR2-KO n=5 for all timepoints).

**Fig. S5.**
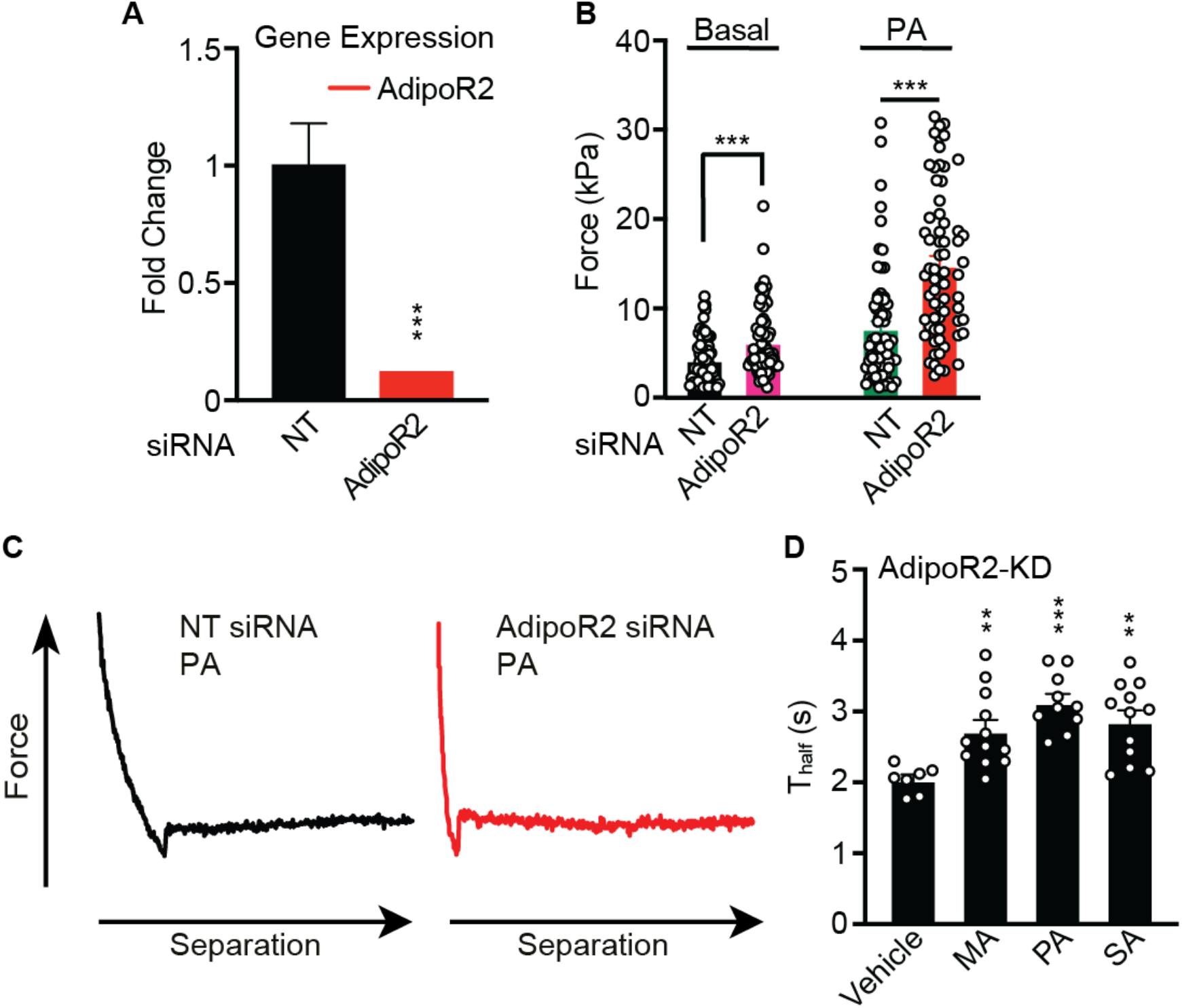
AdipoR2 silencing causes rigidification measured by atomic force microscopy and also alters response to different SFAs. **A**, qPCR results showing the efficiency of AdipoR2 knockdown in HEK293 cells; NT stands for nontarget siRNA. **B**, Membrane deformation measured by atomic force microscopy in HEK293 cells treated with NT and AdipoR2 siRNA under basal conditions or treated with 200 μM PA (NT siRNA and AdipoR2 siRNA Basal, n=66; NT siRNA PA, n=66; AdipoR2 siRNA PA, n=67). **C**, Representative deformation curve for NT and AdipoR2 siRNA-treated cells in the presence of PA **D**, The SFAs myristic acid (MA), PA and stearic acid (SA) all cause membrane rigidification in AdipoR2-knocked down (KD) HEK293 cells as measured using FRAP; the average T_half_ is shown for each condition (n≥7).

**Fig. S6.**
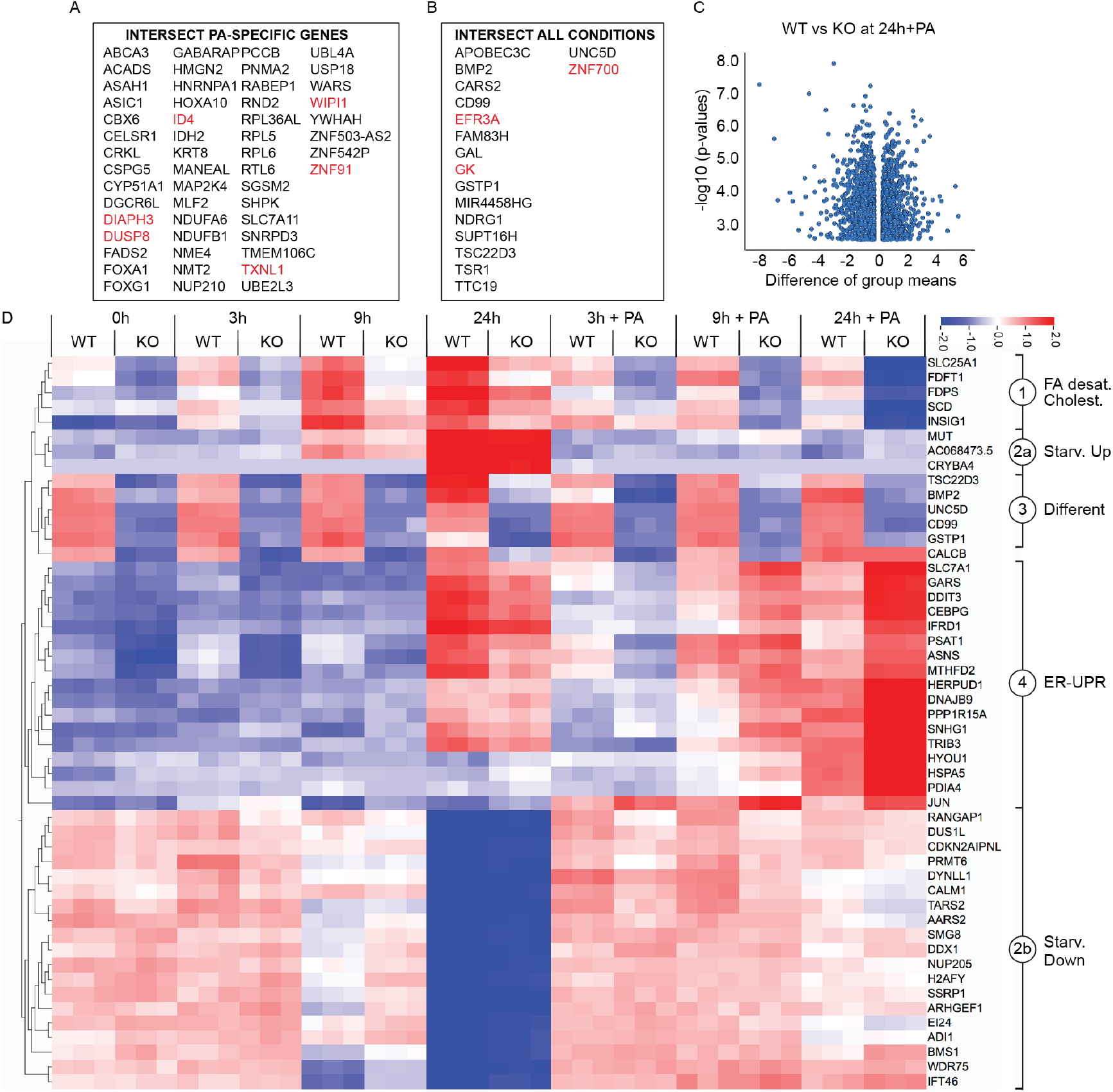
AdipoR2 is required for normal transcriptional response. **A**, List of genes differently expressed at all three time points (3, 9 and 24 h) when the cells are cultivated in the presence of PA (black and red text indicate down- and up-regulated genes, respectively). **B**, List of genes differently expressed at all three time points and under all conditions (full media, serum-free media, and serum-free media + PA) at all three time points (3, 9 and 24 h); black and red text indicate down- and up-regulated genes, respectively. **C**, Volcano plot of the 3050 genes with q<0.05 and at least a 1.2-fold difference between WT and AdipoR2-KO cells after 24 h in serum-free + PA. **D**, Heat map of the top 50 genes showing the most significant variation (p≤4.5e-26) in a multigroup comparison among all treatments. The treatments, performed in triplicates, include: full medium, 3, 9 and 24 h starvation in serum-free medium and 3, 9 and 24 h in serum-free medium containing 200 μM PA for wild-type (WT) and AdipoR2-KO (KO) HEK293 cells. Similarly expressed genes clustered as per the labels on the right-hand side: fatty acid desaturation and cholesterol biosynthesis genes (“FA desat., Cholest.”), genes downregulated or upregulated specifically after 24 h of starvation (“Starv Up” and “Starv. Down”), genes misregulated and related to ER-UPR (“ER-UPR”) and genes likely reflecting the differentiation state (“Different”).

**Fig. S7.**
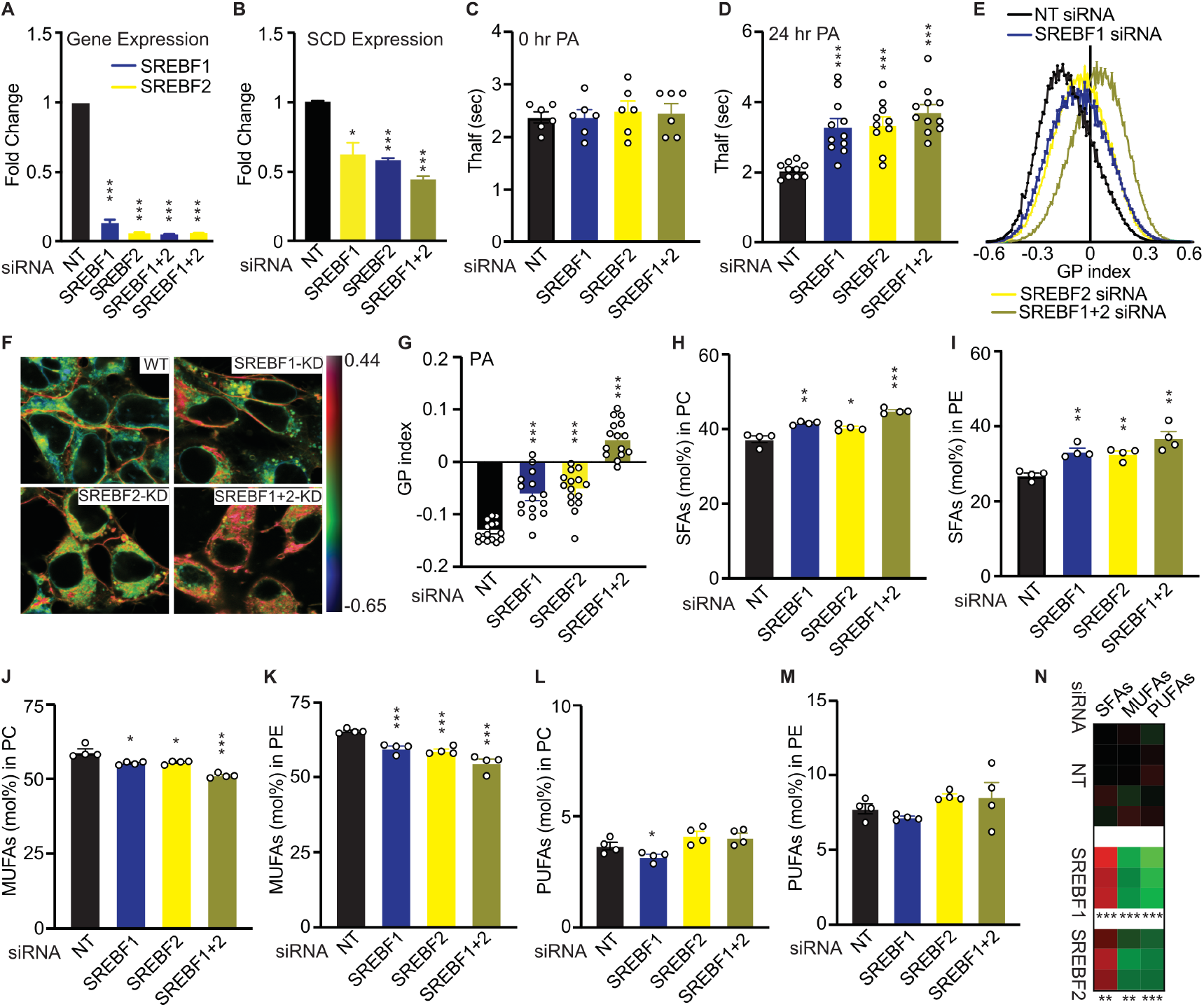
The SREBFs regulate membrane composition and fluidity in HEK293 cells challenged with 200 μM PA. **A**, qPCR results showing the efficiency of different knockdowns in HEK293 cells (n=3 for all treatments). **B**, Relative SCD expression in cells treated with different siRNAs (n=3 for all treatments). **C**,**D** Average T_half_ values comparing cells treated with NT, SREBF1, SREBF2 and SREBF1+2 siRNA after **C**, 0 h PA challenge (n=6 for all treatments) and **D**, 24 h PA challenge (NT siRNA, n=10; SREBF1 siRNA, n=11; SREBF2 siRNA, n=10; SREBF1+2 siRNA, n=11). **E**, Distribution of GP index values for cells treated with SREBF1, SREBF2, SREBF1+2 siRNA and challenged with PA. **F,** Representative pseudocolor images of laurdan dye GP index and **G,** average GP index in HEK293 cells treated with NT, SREBF1, SREBF2 and SREBF1+2 siRNA and challenged with 200 μM PA (n=15 for all treatments). **H-M**, SFA, MUFA and PUFA abundance (mol%) in PCs and PEs in cells treated with different siRNAs and 200 μM PA (n=4 for all treatments). **N**, Relative levels of SFAs, MUFAs and PUFAs in PCs in HUVECs treated with NT, SREBF1 and SREBF2 siRNA and challenged with 200 μM PA for 6 h (NT siRNA, n=5; SREBF1 siRNA, n=3; SREBF1 siRNA, n=3).

**Fig. S8.**
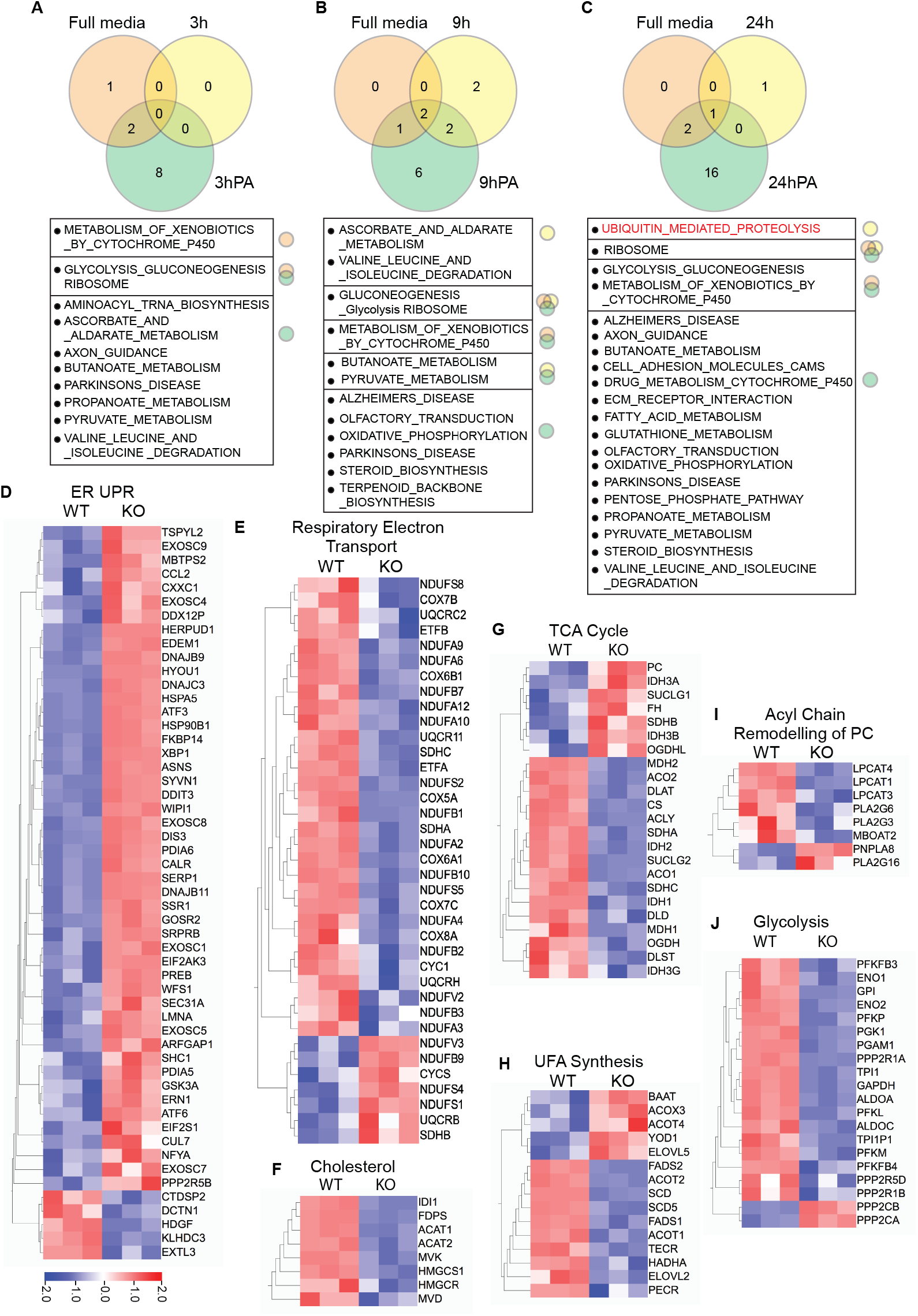
Abnormal transcriptional response in AdipoR2-KO cells challenged with 200 μM PA. **A-C**, Venn diagrams of the KEGG pathways showing significant differences (q<0.05) in a GSEA between WT and AdipoR2-KO cells under each condition (full media, serum-free media, and serum-free media + PA) and time point (3 h, 9 h and 24 h). The affected pathways are listed below each Venn diagram. **D-J**, Heat map comparing the expression of genes from several pathways in WT and AdipoR2-KO cells after 24 h in serum-free media supplemented with 200 μM PA. For each pathway, only genes showing significant differences (p≤0.05) were included in the heat maps. The pathways are: REACTOME Unfolded Protein Response (“ER UPR”), REACTOME Respiratory electron transport (“Respiratory Electron Transport), KEGG Terpenoid Backbone Biosynthesis (“Cholesterol”), REACTOME Respiratory electron transport (“Respiratory Electron Transport), KEGG Citrate Cycle TCA Cycle (“TCA Cycle”), KEGG Biosynthesis of Unsaturated Fatty Acids (“UFA Synthesis”), REACTOME Acyl Chain Remodelling of PC (“Acyl Chain Remodelling of PC”) and REACTOME Glycolysis (“Glycolysis”).

**Fig. S9.**
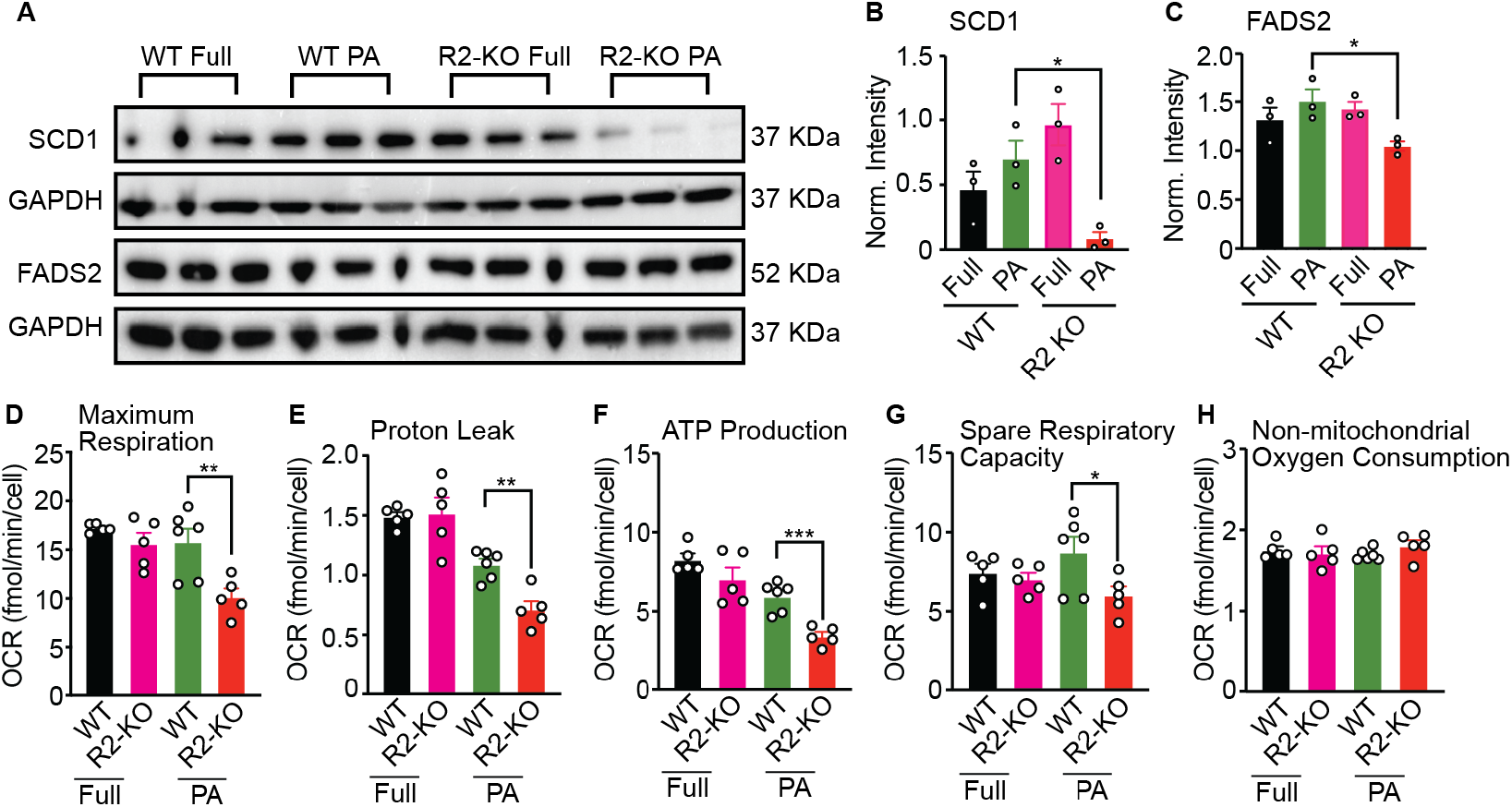
AdipoR2 is required for normal mitochondrial respiration when challenged with 200 μM PA. **A**, Western blot detection of SCD1 and FADS2 in WT and AdipoR2-KO cells in full media or serum-free media + 200 μM PA. **B-C**, GAPDH-based normalized expression of SCD1 and FADS2 from the blot in A (n=3 for all genotypes and conditions). **D-H**, Respiration parameters in WT and AdipoR2-KO cells in full media or serum-free media + 200 μM PA (WT Full, n=5; AdipoR2-KO Full, n=5; WT PA, n=6; AdipoR2-KO PA, n=5).

**Fig. S10.**
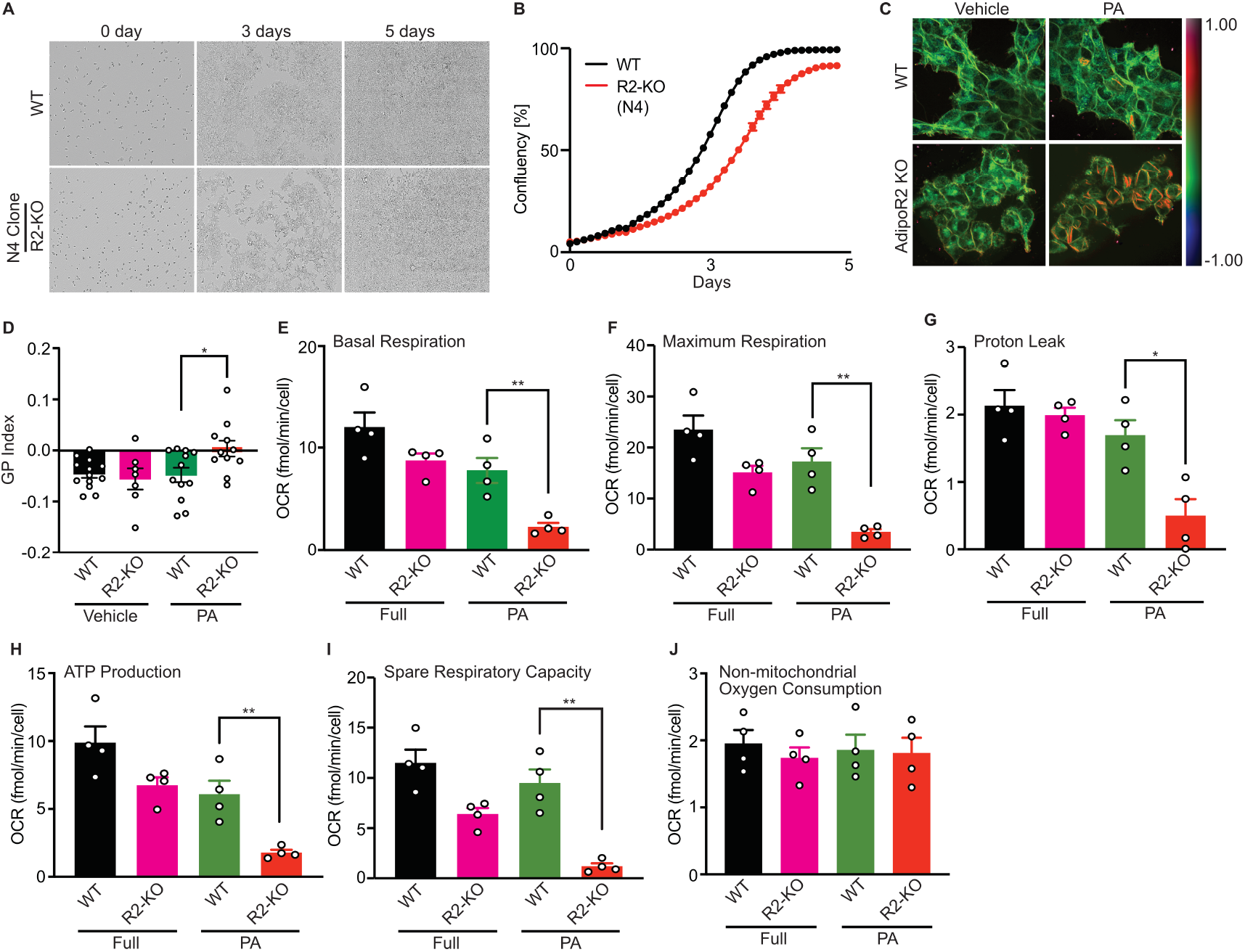
PA intolerance in a HEK293T AdipoR2-KO clone. **A,B,** Growth of the AdipoR2-KO clone N4 is impaired in the presence of 200 μM PA. **C,** Laurdan dye staining shows membrane rigidification in the AdipoR2-KO clone N4 challenged with 200 μM PA. **D,** Average GP index in control (WT) and AdipoR2-KO clone N4 in control media (vehicle) or in the presence of 200 μM PA (WT vehicle n=13, R2-KO vehicle n=7, WT PA n=12, R2-KO PA n=11). **E-J,** Seahorse measurements show that several mitochondrial respiration parameters are impaired in the AdipoR2-KO clone N4 challenged with 200 μM PA (n=4 for each condition).

**Fig. S11.**
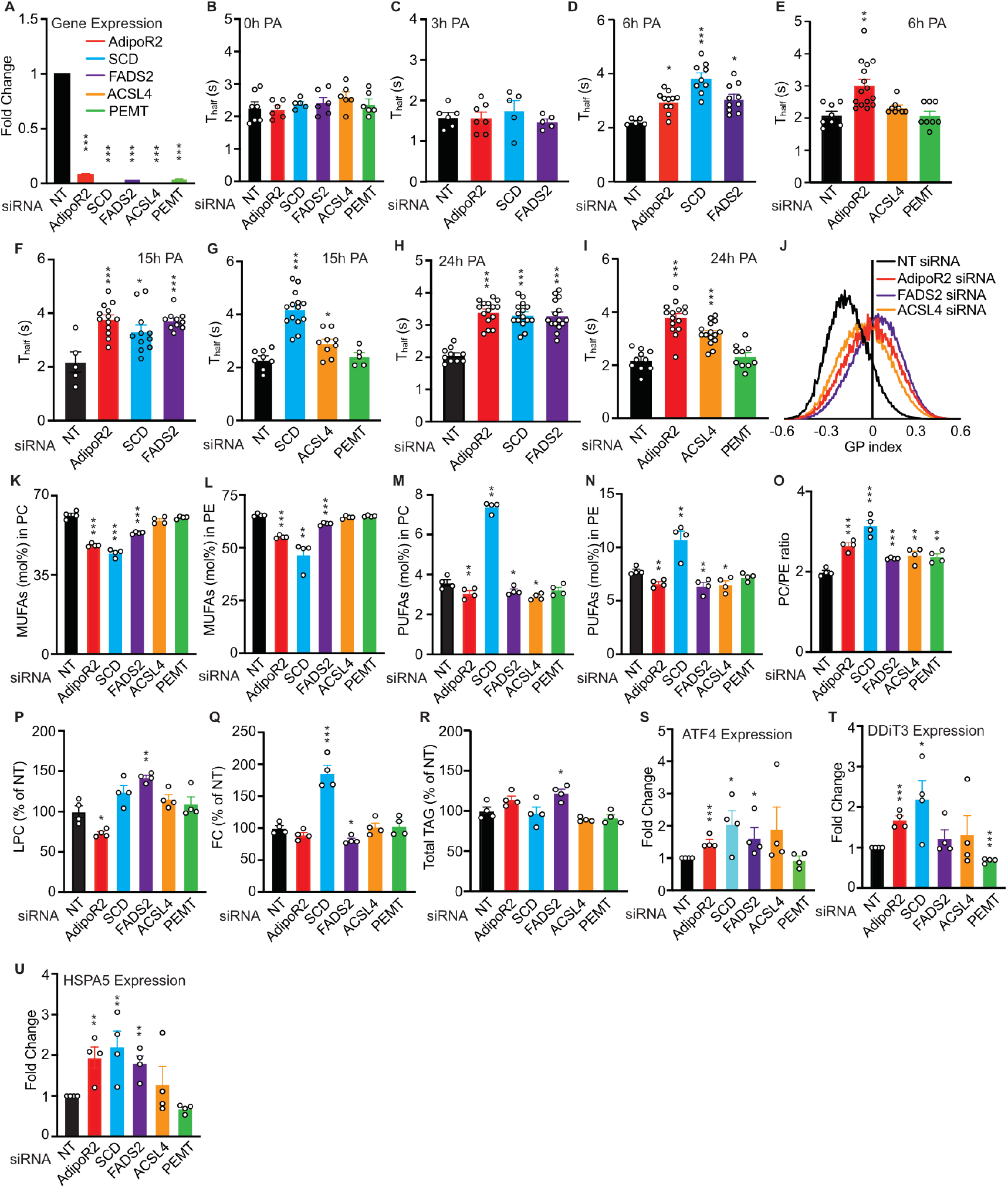
Importance of AdipoR2 to regulate membrane homeostasis. **A**, qPCR results showing the efficiency of different knockdowns. **B**, Average T_half_ values from FRAP experiments comparing cells treated with NT, AdipoR2, SCD, FADS2, ACSL4 and PEMT siRNA after 0 h PA (NT siRNA, n=7; AdipoR2 siRNA, n=6; SCD siRNA, n=5; FADS2 siRNA, n=6; ACSL4 siRNA, n=6; PEMT siRNA, n=6). **C**, Average T_half_ values comparing cells treated with NT, AdipoR2, SCD, FADS2, siRNA after 3 h 200 μM PA (NT siRNA, n=6; AdipoR2 siRNA, n=7; SCD siRNA, n=5; FADS2 siRNA, n=5). **D**, **E**, Average T_half_ values comparing cells treated with NT, AdipoR2, SCD, FADS2, ACSL4 and PEMT siRNA after 6 h PA (NT siRNA, n=8; AdipoR2 siRNA, n=15; SCD siRNA, n=9; FADS2 siRNA, n=10; ACSL4 siRNA, n=8; PEMT siRNA, n=8). **F**, **G**, Average T_half_ values comparing cells treated with NT, AdipoR2, SCD, FADS2, ACSL4 and PEMT siRNA after 15 h PA (NT siRNA, n=5; AdipoR2 siRNA, n=13; SCD siRNA, n=11; FADS2 siRNA, n=10; ACSL4 siRNA, n=9; PEMT siRNA, n=5). **H**, **I**, Average T_half_ values comparing cells treated with NT, AdipoR2, SCD, FADS2, ACSL4 and PEMT siRNA after 24 h 200 μM PA c (NT siRNA, n=10; AdipoR2 siRNA, n=15; SCD siRNA, n=15; FADS2 siRNA, n=15; ACSL4 siRNA, n=15; PEMT siRNA, n=10). **J**, Distribution of GP index values for cells treated with NT, AdipoR2, SCD, FADS2 siRNA and challenged with PA. **K-N**, MUFAs and PUFAs abundance (mol%) in PCs and PEs in cells treated with different siRNAs and PA (n=4 for all treatments). **O-R**, PC/PE ratio, relative lysophosphatidylcholine (LPC), free cholesterol (FC) and total TAGs in cells treated with different siRNAs and PA (n=4 for all treatments). **S-U**, Relative gene expression for ATF4, DDiT3 and HSPA5 in cells treated different siRNAs and PA (n=4 for all treatments).

## Notes

### Competing Interest Statement

The authors have declared no competing interest.

